# Glycemic challenge is associated with the rapid cellular activation of the locus ceruleus and nucleus of solitary tract: Circumscribed spatial analysis of phosphorylated MAP kinase immunoreactivity

**DOI:** 10.1101/2022.08.30.504809

**Authors:** Geronimo P. Tapia, Lindsay J. Agostinelli, Sarah D. Chenausky, Jessica V. Salcido Padilla, Vanessa I. Navarro, Amy Alagh, Gabriel Si, Richard H. Thompson, Sivasai Balivada, Arshad M. Khan

## Abstract

Rodent studies indicate that impaired glucose utilization or hypoglycemia is associated with cellular activation of neurons in the *medulla* (*Winslow, 1733*) (MY) believed to control feeding behavior and glucose counterregulation. However, such activation has been tracked primarily within hours of the challenge, rather than sooner, and has been poorly mapped within standardized brain atlases. Here, we report that within 15 min of receiving 2-deoxy-D-glucose (2-DG; 250 mg/kg, i.v.), which can trigger glucoprivic feeding behavior, marked elevations were observed in the numbers of *rhombic brain* (*His, 1893*) (RB) neuronal cell profiles immunoreactive for the cellular activation marker(s), phosphorylated p44/42 MAP kinases (phospho-ERK1/2), some of which were also catecholaminergic. We mapped their distributions within an open-access rat brain atlas and found that 2-DG-treated rats (compared to their saline-treated controls) displayed greater numbers of phospho-ERK1/2^+^ neurons in the *locus ceruleus* (*Wenzel & Wenzel, 1812*) (LC) and the *nucleus of solitary tract (>1840)* (NTS). Thus, 2-DG-activation of certain RB neurons is more rapid than perhaps previously realized, engaging neurons that serve multiple functional systems and are of varying cellular phenotypes. Mapping these populations within standardized brain atlas maps streamlines their targeting and/or comparable mapping in preclinical rodent models of disease.

## 1. Introduction

Glucose is the essential metabolic fuel source for the brain, with a constant and adequate supply needed for proper function and survival. Iatrogenic hypoglycemia is a formidable clinical problem for patients suffering from diabetes mellitus [1], and despite steady improvements in long-term diabetes management [2], remains a major limiting factor in such management for both Type I and Type II diabetics [1, 3–5]. Glucose counterregulatory mechanisms defend the body against precipitous drops in blood sugar by increasing glucagon and reducing insulin secretion from pancreatic α and β cells, respectively; and by increasing epinephrine release from the adrenal medulla [6]. In Type I and advanced (end-stage) Type II diabetes, these defenses are impaired, thereby compromising the body’s ability to maintain glucose homeostasis. Such impairment becomes exacerbated during hypoglycemia-associated autonomic failure (HAAF), in which antecedent iatrogenic hypoglycemia causes defective glucose counterregulation by blunting plasma epinephrine responses to a given level of subsequent hypoglycemia — all in the face of impaired glucagon release and insulin reductions [7]. In addition to defective glucose counterregulation, HAAF is often associated with hypoglycemia unawareness, in which “cognitive function is so disturbed that the patient can become drowsy, uncoordinated, confused or even comatose” [8].

Human and animal models for HAAF exist but need refinement [9], partly because not all of the fundamental mechanisms underlying counterregulation have been elucidated [10]. Although sensors in the periphery (e.g., portal vein [11]) contribute to counterregulatory responses to glycemic challenges, there is a growing understanding that brain mechanisms also contribute to these responses [12–15]. Therefore, there have been intensive efforts to understand how the *central nervous system* (*Carus, 1814*) [16] (CNS) (*see Section 2.1 for naming conventions used in this study*) senses circulating levels of glucose and initiates counterregulatory responses, and how CNS impairments contribute to defective glucose counterregulation [17–20]. A compelling body of evidence has established a critical role for various *rhombic brain* (*His, 1893*) [21] (RB) regions in glucosensing and counterregulatory responses to glycemic challenges [22–36], with much of this work focused on the role of catecholaminergic neurons in the *medulla* (*Winslow, 1733*) [37] (MY) in these functions [17, 38–42]. Certain neuronal populations in the *hypothalamus* (*Kuhlenbeck, 1927*) [54] (HY) also appear to be glucosensing and/or critical for counterregulation [55–65], and neural circuits linking the RB to the HY help trigger feeding and hypothalamo-pituitary-adrenal (HPA) axis responses to glycemic challenges [15, 66–73]. Moreover, peripheral glucosensing mechanisms can be linked by neural circuits to RB regions controlling counterregulatory responses [18, 19, 74–76].

Despite these advances in our understanding of RB glucosensing and counterregulation, two points regarding the spatiotemporal dynamics of these responses remain unclear, both of which relate to the aims of the present study. First, in terms of temporal dynamics, it is not fully understood how quickly RB neuronal networks track changes in glycemic status *in vivo*. Immunodetection of the transcription factor, Fos, has been used to track RB neuronal activation associated with glycemic challenge [30, 39, 42, 53, 75], but its protein expression typically peaks within 1–3 hours [77]. Thus, it is difficult to attribute its delayed expression to changes in glycemic status per se rather than to secondary effects set in motion by such changes. Ideally, an anatomically non-labile biomarker of an “activated” neuron should be induced at a time more proximate to the stimulus event (discussed in [79]). In this study, the phosphorylated forms of p44/42 MAP kinases were used to rapidly track glycemic challenge, a strategy first employed by the Watts laboratory [66–69].

Second, in terms of spatial dynamics, the locations of RB neuronal populations that are activated following glycemic challenge have not been charted precisely within standardized atlases of the brain, rendering their reproducible experimental targeting and documentation more difficult (discussed in [80]). Nor is it clear if RB neuronal populations more traditionally studied in the context of arousal rather than hormonal counterregulation are also activated rapidly by glycemic challenge. This knowledge could help us understand if such regions might play a role in producing the cognitive perceptions of hypoglycemic events, the absence of which could contribute to hypoglycemia unawareness in diabetic individuals [8, 81]. Here, we build upon previous work [68] to map RB neuronal activation associated with glycemic challenge to an open-access rat brain atlas [82] and identify the *locus ceruleus* (*Wenzel & Wenzel, 1812*) [83], a key arousal-promoting RB structure, as a region displaying rapid activation to glycemic challenge, alongside other RB structures more traditionally understood to serve in networks for glucosensing and counterregulation. Taken together, the results of this work, portions of which have been presented in preliminary form [85–90], provide evidence that RB neuronal populations implicated in glucosensing, counterregulation and arousal can track the onset of glycemic challenge within 15 minutes. Importantly, our work provides two sets of novel results addressing the aims of our study: (1) demonstration of rapid temporal activation in the RB following glycemic challenge; and (2) precise spatial mapping of these activation patterns using an open-access brain atlas. The cellular activation patterns mapped in this study are a useful starting point towards the development of global CNS cellular activation maps for glycemic challenges.

## 2. Materials and Methods

### 2.1 Neuroanatomical naming conventions used in this study

In this study, as part of our ongoing effort to encourage the use of standardized nomenclature to enable the seamless registration of datasets across laboratories (and datasets within our own laboratory), we use *standard terms* for brain structures as proposed by Swanson [82, 91]. These terms are listed in italics together with the associated citation that first uses the term as defined. Note that the author-date citation is *part of the standard term*, and these nomenclature-embedded citations are also included in the references list. If a definitive assignment of priority for the term was not possible, it was assigned by Swanson the citation “(>*1840*)”; that is, “defined sometime after the year 1840″. Refer to Swanson [82, 91] for further details regarding this standard nomenclature system.

### 2.2 Subjects and surgical procedures

All procedures were approved by the Institutional Animal Care and Use Committee of the University of Southern California (Protocol code #11239). Adult male Sprague-Dawley rats (Harlan Labs, Indianapolis, IN, USA; n = 12; body weights (BW) 258–270 g on the day of surgery) were single-housed and kept on a 12h:12h light:dark cycle (lights on: 6:00 a.m.) with *ad libitum* access to standard laboratory animal chow and water. On the day of surgery, each rat was anesthetized with a cocktail of 50% ketamine (100 mg/ml), 25% xylazine (100 mg/ ml), and 10% acepromazine (10 mg/ml) dissolved in 0.9% sterile saline; delivered i.m. at 0.1 ml of cocktail/100 g BW). The animal was judged to be at the appropriate surgical plane of anesthesia by tracking expected reductions in respiratory rate following anesthesia, as well as the absence of a reflex response to a brief tail or paw pinch.

Intra-atrial catheterizations by way of the jugular vein, under aseptic surgical conditions, were performed as previously reported [66, 92], with additional details noted in the *Supplemental Methods*. Post-operative care consisted of administration of flunixin meglumine analgesic (0.25 mg/100 g BW, i.m.) and gentamicin antibiotic (0.2–0.4 mg/100 g BW, i.m.), and the careful daily monitoring and gentle handling of each animal throughout the course of the 3–6-day recovery period, during which time each animal was weighed to determine whether they reached pre-surgical body weight. During this recovery period, their newly-installed catheters were also inspected and flushed daily with heparinized saline to ensure their patency.

### 2.3 In vivo experimental procedures

On the day of the experiment, during the early portion of the light phase, the animals received an injection, through the catheter, of 0.9% sterile saline as a vehicle control (n = 7), or 250 mg/kg of 2-deoxy-D-glucose (2-DG; catalog #D6134-5G; MilliporeSigma, Inc., Burlington, MA, USA) dissolved in the vehicle (n = 5). This intravenous procedure was intended to eliminate pain-related neuronal activation that would otherwise occur by administering the treatment through intraperitoneal injection. Approximately fifteen minutes after receiving saline or 2-DG, subjects received an intravenous infusion of ~1.0 ml sodium pentobarbital anesthesia (50 mg/kg) over a 30-sec period. Deeply sedated, the subjects were relocated to a fume hood, under which they were perfused transcardially, first with 0.01 M sodium phosphate-buffered saline and then with freshly depolymerized 4% (w/v) *p*-formaldehyde, buffered to pH 9.5 using sodium borate. The fixed brain was removed from the cranium, placed overnight on a shaker at 4°C in 4% (w/v) *p*-formaldehyde containing 12% (w/v) sucrose, blocked into forebrain and hindbrain segments, and then frozen using hexane supercooled on a bed of powdered dry ice. The frozen blocks were removed from the hexane bath and stored in vials at −80°C until sectioning.

### 2.4 Tissue processing and immunocytochemistry

#### 2.4.1 Histological processing of brain tissue

Rhombic brain (His, 1893) [21] (RB) blocks were removed from −80°C and mounted onto the freezing stage of a Reichert sliding microtome and cut into 20-μm-thick coronal sections. Sections kept for histochemical staining were collected as eight 1-in-8 series in tissue culture wells filled with cryoprotectant (50% phosphate buffer, 30% ethylene glycol, 20% glycerol; pH 7.4) and stored at –20°C until further processing. In most cases, the fourth tissue series collected was reserved for preparing a cytoarchitectural *Reference (R) series* using Nissl-staining (*see Section 2.5*), while the remaining tissue series were reserved for immunohistochemical analyses. The *R series* for any one subject is defined as the tissue series from which we used cytoarchitectural information to identify the boundary conditions for mapping the atlas *gray matter regions (Swanson & Bota, 2010)* of our immunoreactive neuronal populations, which were visualized in adjacent tissue series from the same subject.

#### 2.4.2 Immunohistochemistry

##### 2.4.2.1. Main procedures

Using Tris-buffered saline (TBS) (0.05 M; pH 7.4 at room temperature), sections were rinsed of cryoprotectant (5 rinses for 5 minutes each; 5 × 5) and were then incubated in blocking solution consisting of TBS containing 2% (v/v) normal donkey serum (Catalog # S30–100ML; EMD-Millipore), and 0.1% (v/v) Triton X-100 for ~1.5–2.0 h at room temperature. The tissue was reacted with a cocktail consisting of blocking solution together with primary antibodies targeting dopamine-β-hydroxylase ([EC 1.14.17.1](https://iubmb.qmul.ac.uk/enzyme/EC1/14/17/1.html)), choline acetyltransferase ([EC 2.3.1.6](https://iubmb.qmul.ac.uk/enzyme/EC2/3/1/6.html)), or the phosphorylated forms of ERKs 1 and 2 ([EC 2.7.11.24](https://iubmb.qmul.ac.uk/enzyme/EC2/7/11/24.html)) (*see* **Table 1***for details*). After another 5 × 5 TBS rinse, the sections were reacted with the appropriate secondary cocktail (again with blocking solution), followed by a conjugate reagent **(Table 1)**. The tissue underwent a final 5 × 5 wash series before being mounted on Superfrost™ Plus slides, which were coverslipped with sodium bicarbonate-buffered glycerol (pH 8.6 at room temperature), sealed with clear nail polish, stored at 4°C, and protected from ambient light until further analysis.

**Table 1.**
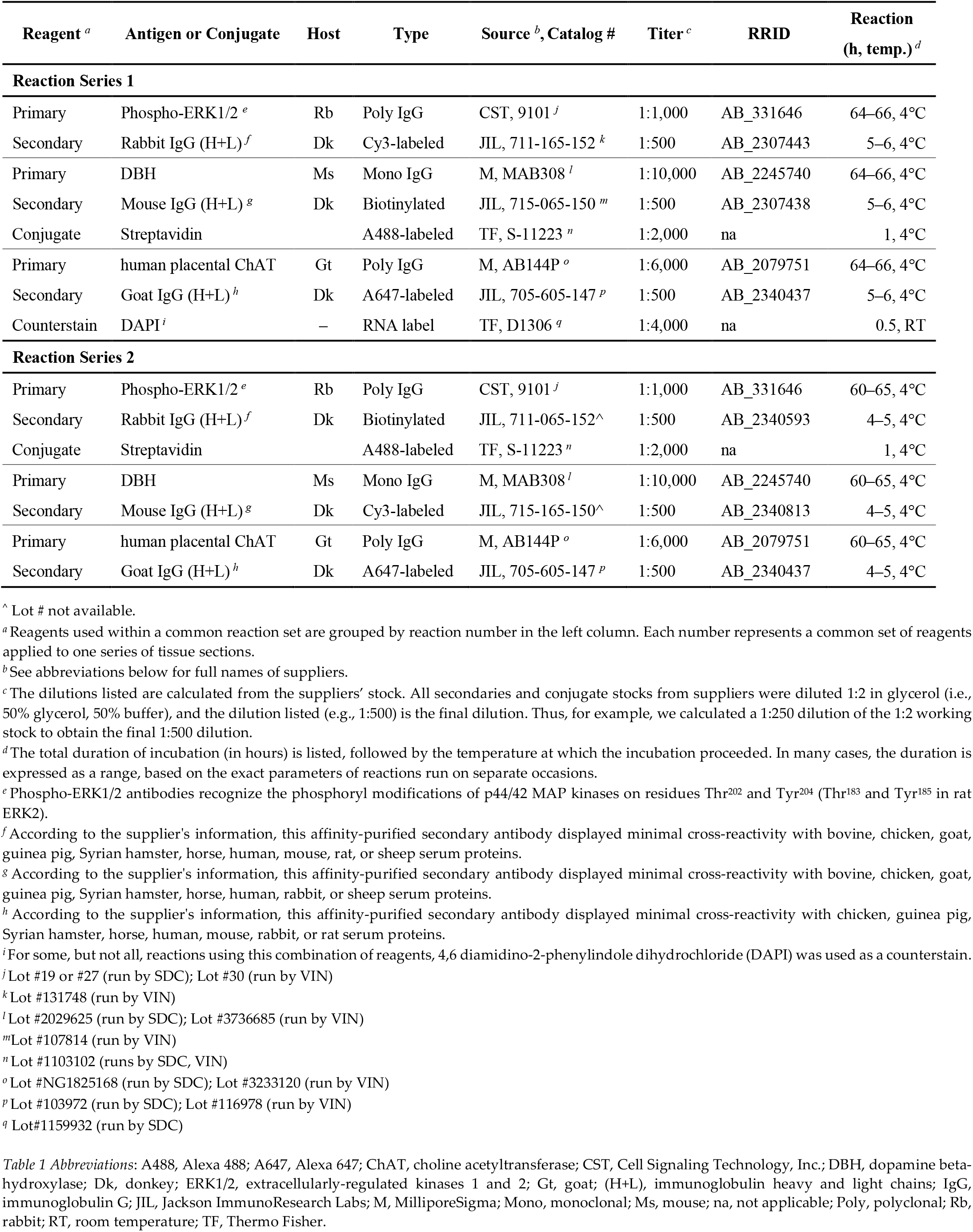
Antibody combinations

| Reagent <sup>a</sup> | Antigen or Conjugate | Host | Type | Source <sup>b</sup>, Catalog # | Titer <sup>c</sup> | RRID | Reaction (h, temp.) <sup>d</sup> |
| --- | --- | --- | --- | --- | --- | --- | --- |
| **Reaction Series 1** |  |  |  |  |  |  |  |
| Primary | Phospho-ERK1/2 <sup>e</sup> | Rb | Poly IgG | CST, 9101 <sup>j</sup> | 1:1,000 | AB_331646 | 64–66, 4°C |
| Secondary | Rabbit IgG (H+L) <sup>f</sup> | Dk | Cy3-labeled | JIL, 711-165-152 <sup>k</sup> | 1:500 | AB_2307443 | 5–6, 4°C |
| Primary | DBH | Ms | Mono IgG | M, MAB308 <sup>l</sup> | 1:10,000 | AB_2245740 | 64–66, 4°C |
| Secondary | Mouse IgG (H+L) <sup>g</sup> | Dk | Biotinylated | JIL, 715-065-150 <sup>m</sup> | 1:500 | AB_2307438 | 5–6, 4°C |
| Conjugate | Streptavidin |  | A488-labeled | TF, S-11223 <sup>n</sup> | 1:2,000 | na | 1, 4°C |
| Primary | human placental ChAT | Gt | Poly IgG | M, AB144P <sup>o</sup> | 1:6,000 | AB_2079751 | 64–66, 4°C |
| Secondary | Goat IgG (H+L) <sup>h</sup> | Dk | A647-labeled | JIL, 705-605-147 <sup>p</sup> | 1:500 | AB_2340437 | 5–6, 4°C |
| Counterstain | DAPI <sup>i</sup> | – | RNA label | TF, D1306 <sup>q</sup> | 1:4,000 | na | 0.5, RT |
| **Reaction Series 2** |  |  |  |  |  |  |  |
| Primary | Phospho-ERK1/2 <sup>e</sup> | Rb | Poly IgG | CST, 9101 <sup>j</sup> | 1:1,000 | AB_331646 | 60–65, 4°C |
| Secondary | Rabbit IgG (H+L) <sup>f</sup> | Dk | Biotinylated | JIL, 711-065-152 <sup>^</sup> | 1:500 | AB_2340593 | 4–5, 4°C |
| Conjugate | Streptavidin |  | A488-labeled | TF, S-11223 <sup>n</sup> | 1:2,000 | na | 1, 4°C |
| Primary | DBH | Ms | Mono IgG | M, MAB308 <sup>l</sup> | 1:10,000 | AB_2245740 | 60–65, 4°C |
| Secondary | Mouse IgG (H+L) <sup>g</sup> | Dk | Cy3-labeled | JIL, 715-165-150 <sup>^</sup> | 1:500 | AB_2340813 | 4–5, 4°C |
| Primary | human placental ChAT | Gt | Poly IgG | M, AB144P <sup>o</sup> | 1:6,000 | AB_2079751 | 60–65, 4°C |
| Secondary | Goat IgG (H+L) <sup>h</sup> | Dk | A647-labeled | JIL, 705-605-147 <sup>p</sup> | 1:500 | AB_2340437 | 4–5, 4°C |

<sup>^</sup> Lot # not available.
<sup>a</sup> Reagents used within a common reaction set are grouped by reaction number in the left column. Each number represents a common set of reagents applied to one series of tissue sections.
<sup>b</sup> See abbreviations below for full names of suppliers.
<sup>c</sup> The dilutions listed are calculated from the suppliers' stock. All secondaries and conjugate stocks from suppliers were diluted 1:2 in glycerol (i.e., 50% glycerol, 50% buffer), and the dilution listed (e.g., 1:500) is the final dilution. Thus, for example, we calculated a 1:250 dilution of the 1:2 working stock to obtain the final 1:500 dilution.
<sup>d</sup> The total duration of incubation (in hours) is listed, followed by the temperature at which the incubation proceeded. In many cases, the duration is expressed as a range, based on the exact parameters of reactions run on separate occasions.
<sup>e</sup> Phospho-ERK1/2 antibodies recognize the phosphoryl modifications of p44/42 MAP kinases on residues Thr<sup>202</sup> and Tyr<sup>204</sup> (Thr<sup>183</sup> and Tyr<sup>185</sup> in rat ERK2).
<sup>f</sup> According to the supplier's information, this affinity-purified secondary antibody displayed minimal cross-reactivity with bovine, chicken, goat, guinea pig, Syrian hamster, horse, human, mouse, rat, or sheep serum proteins.
<sup>g</sup> According to the supplier's information, this affinity-purified secondary antibody displayed minimal cross-reactivity with bovine, chicken, goat, guinea pig, Syrian hamster, horse, human, rabbit, or sheep serum proteins.
<sup>h</sup> According to the supplier's information, this affinity-purified secondary antibody displayed minimal cross-reactivity with chicken, guinea pig, Syrian hamster, horse, human, mouse, rabbit, or rat serum proteins.
<sup>i</sup> For some, but not all, reactions using this combination of reagents, 4,6 diamidino-2-phenylindole dihydrochloride (DAPI) was used as a counterstain.
<sup>j</sup> Lot #19 or #27 (run by SDC); Lot #30 (run by VIN)
<sup>k</sup> Lot #131748 (run by VIN)
<sup>l</sup> Lot #2029625 (run by SDC); Lot #3736685 (run by VIN)
<sup>m</sup> Lot #107814 (run by VIN)

##### 2.4.2.2 Antibody information and immunohistochemical controls. Please refer to the *Supplemental Methods* for details

### 2.5 Nissl staining of the Reference (R) tissue series

The *R series* of tissue sections (adjacent to those that underwent immunohistochemical staining) was washed 5 × 5 in TBS and mounted onto gelatin-coated slides, dehydrated in ascending concentrations of ethanol (50%, 70%, 95%, 100%) and defatted with xylene. Sections were then rehydrated and stained with a 0.25% w/v thionine solution (sold as “thionin acetate” by Sigma-Aldrich; MW 287.34 g/mol; cat #T7029; ([93], p. 151); [94]). A 0.4% glacial acetic acid solution was used to remove excess thionine and the tissue slides were dehydrated in ascending concentrations of ethanol solutions (50%, 70%, 95%, 100% × 3), and then cleared in mixed-isomer xylenes before being coverslipped using DPX mountant for bright-field microscopic observation (*Section 2.6.1*).

### 2.6 Microscopy and photomicrography

#### 2.6.1 Bright-field microscopy

A total of 93 *R series* (Nissl-stained) tissue sections, for a subset of saline- and 2-DG-treated subjects (n = 2 and 3, respectively), were photographed under bright-field illumination using an Olympus BX63 light microscope with an X-Y-Z motorized stage and an attached DP74 color CMOS camera (cooled, 20.8 MP pixel-shift, 60 fps) (Olympus America, Inc., Waltham, MA, USA). Image acquisition was carried out using cellSense™ Dimension software (Version 2.3; Olympus) installed on a Hewlett Packard PC workstation. The software was used to drive the motorized stage and generate tiled and stitched wide-field mosaic photomicrographs of the entire tissue section using a ×10 magnification objective lens (Olympus UPlanSApo, N.A. 0.40, FN 26.5 mm). Stitching was performed with a 15% overlap of individual images.

#### 2.6.2 Epifluorescence and dark-field microscopy

Selected tissue sections from subjects receiving saline or 2-DG were photographed using epifluorescence illumination. Identifying information was first obscured on all tissue slides by a team member (AMK), and the sections were then photographed by another team member (GPT) who was thus blind to the treatment groups. Photography was carried out using an Axio Imager M.2 upright epifluorescence microscope (Carl Zeiss, Inc., Thornwood, NY, USA) equipped with an X-Y-Z motorized stage and a 100 W halogen light source. The appropriate filter cubes were used to detect the immunofluorescence of the sections in separate non-interfering channels, which was captured by a EXi Blue monochrome camera (Teledyne QImaging, Inc., Surrey, British Columbia, Canada) driven by Volocity software (versions 6.0.0–6.1.1; Quorum Technologies, Puslinch, Ontario, Canada) installed on a Mac Pro computer (Apple Corp., Cupertino, CA, USA). Single fields of view and ×20-magnification (Plan-Apochromat objective lens; N.A., 0.8) wide-field mosaics of individual fields of view (stitched with 20–40% overlap) were acquired within Volocity and exported as uncompressed TIFF files. These photomicrographs were also exported as compressed JPEG files for conference presentation of preliminary results [86, 87]. ROIs were photographed for each subject from the left and right sides of the *rhombic brain (His, 1893)* [21] (RB), and included the *locus ceruleus (Wenzel & Wenzel, 1912)* [83], *area postrema (>1840)*, *nucleus of the solitary tract, medial part (>1840)*, *dorsal motor nucleus of vagus nerve (>1840)*, and *hypoglossal nucleus (>1840)*. Care was taken to ensure that all photomicrographs were captured under similar illumination conditions. For certain sections where the companion *R series* tissue was damaged or otherwise unavailable for reference, a photomicrograph was taken – under dark-field illumination – of the fluorescence tissue series in order to obtain fiducial landmarks to help delineate cytoarchitectural boundaries and to determine the atlas level assignment of the tissue section.

#### 2.6.3 Confocal microscopy

For selected *rhombic brain (His, 1893)* [21] (RB) tissue sections, additional imaging was performed using a Zeiss LSM 700 laser scanning confocal system connected to a Zeiss Observer.Z1 inverted fluorescence microscope (Carl Zeiss, Inc., Thornwood, NY, USA), located in the Cellular Characterization and Biorepository Core Facility of the Border Biomedical Research Center at UT El Paso. The system is equipped with four solid-state lasers, including those at 488-nm, 555-nm, and 639-nm wavelengths, which were used to detect Alexa 488, Cy3, and Alexa 647 fluorophore signals, respectively. The microscope was driven by Zen 2009 software (release version 6.0 SP2; version 6.0.0.303; Carl Zeiss Microimaging GmbH).

Once it was determined that the signal was within the detector’s dynamic range (see *Supplemental Methods* for further details), sequential unidirectional scans of the tissue were taken using Plan-Neofluar objective lenses of ×10 (N.A., 0.3; Ph1), ×20 (N.A., 0.5), or ×40 (N.A., 1.3; oil) magnification. Pinhole settings were kept at or close to 1 Airy unit [95] for each channel, defined as the width of the point-spread function for that channel; however, for images used for co-localization analysis, the pinholes were adjusted above or below the 1 Airy unit setting so that the widths of the optical sections for each channel were matched in the axial (*z*) dimension to ensure accurate comparison of signals [96]. With a few exceptions, frame sizes were set to 2048 × 2048 pixels. Image sizes were 618.9 μm × 618.9 μm (pixel size = 0.30 μm). Scanning was performed at a slow scan speed (setting: 4) with 16 frames averaged over a scan time of ~36 minutes (~12 min/channel). Scans were exported as full-resolution uncompressed TIFF files and imported onto artboards in Adobe Illustrator CC (version 25.0; Adobe Systems, Inc., San Jose, CA, USA) to prepare figures for this study.

#### 2.6.4 Checks for fluorescence crosstalk

A number of quality-control checks, which are also recommended as community standards (*e.g*., [96, 97]) were performed to evaluate crosstalk (bleed-through) of the fluorescence signals we detected in this study. These included evaluations of crosstalk at the level of fluorescence labeling, epifluorescence imaging, and confocal imaging (see *Supplemental Methods* for details).

### 2.7 Spatial analyses and tabulation

#### 2.7.1 Sampled regions

For this initial characterization of immunofluorescent labeling patterns in saline- and 2-DG-treated subjects, we sampled selected regions in the *rhombic brain (His, 1893)* [21] (RB) at three selected rostrocaudal levels **(Figure 1)**. Specifically, the tissue sections we examined were sampled at inferred anteroposterior stereotaxic coordinates of –9.80 mm from Bregma (β) (corresponding to Level 51 of BM4.0), –13.44 mm from β (Level 67), and –13.76 from β (Level 69). Our analyses included observations of immunofluorescent patterns in portions of the *locus ceruleus (Wenzel & Wenzel, 1912)* [83] (LC), *area postrema (>1840)*, *nucleus of solitary tract (>1840)* (NTS), *dorsal motor nucleus of vagus nerve (>1840)* (DMX), and *hypoglossal nucleus (>1840)* (XII).

**Figure 1.**
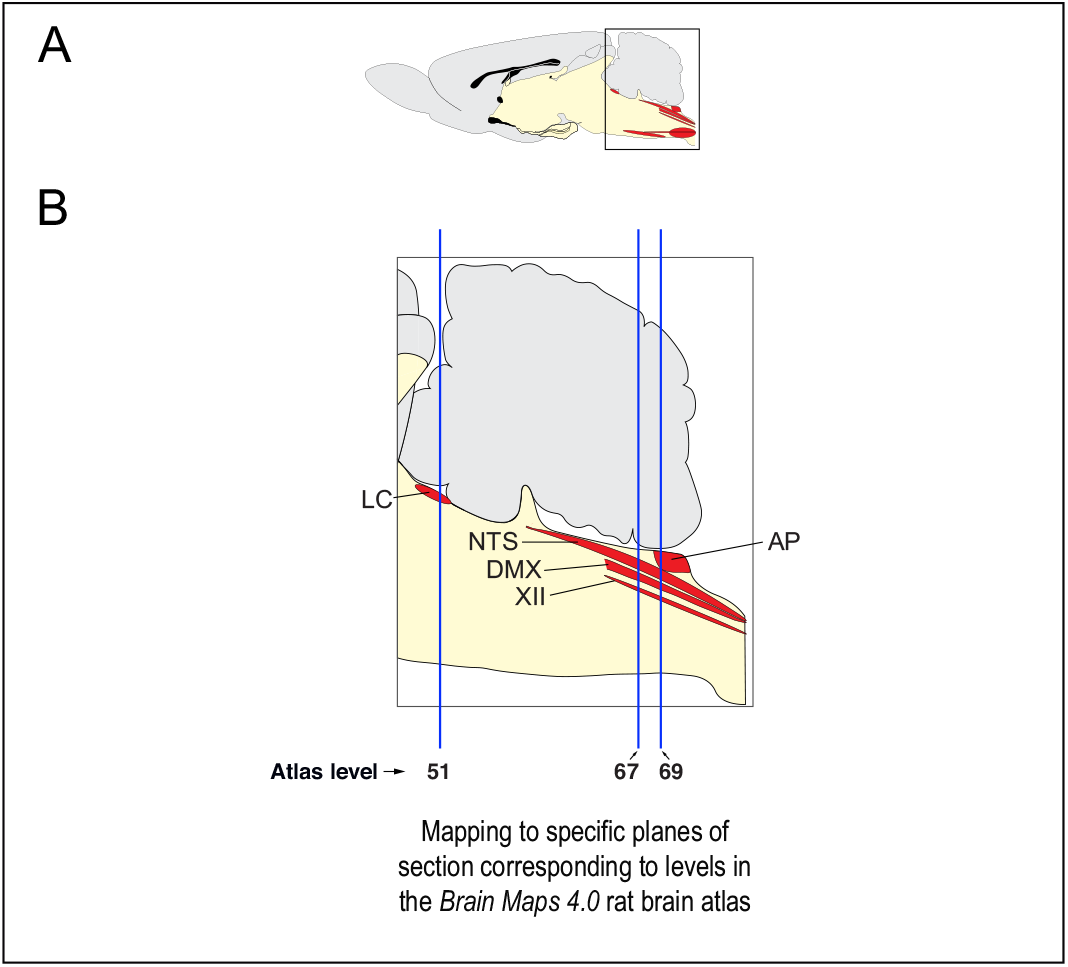
Overview of tissue sampling employed for the experimental dataset. **(A)** A sagittal-plane view of the rat brain, adapted from the *Brain Maps 4.0 (BM4.0)* open-access rat brain atlas [82], which is the spatial model we used in this study. **(B)** An enlargement is presented of the boxed outline shown in **(A)**, which provides details concerning the portions of the *rhombic brain (His, 1893)* [21] (RB) we studied (*shaded in red*). These include the *locus ceruleus (Wenzel & Wenzel, 1912)* [83] (LC), *area postrema (>1840)* (AP), *nucleus of solitary tract (>1840)* (NTS), *dorsal motor nucleus of vagus nerve (>1840)* (DMX), and *hypoglossal nucleus (>1840)* (XII). Portions of these structures were identified in tissue sections as closely registered to specific atlas levels from the *BM4.0* reference (*shown as vertical blue lines*). See the methods section for details.

#### 2.7.2 Plane-of-section analysis, regional parcellations and atlas level assignments

To determine whether the tissue sections were at the rostrocaudal locations that corresponded to the atlas levels we aimed to sample, we performed a plane-of-section analysis for each tissue section collected. We have briefly reported on this type of analysis for Fos-immunoreactive tissue under conditions of fasting and re-feeding [98], and note here additional details germane to the present study (for a more thorough review of this topic, please consult Simmons & Swanson [99]).

First, all *R series* tissue was evaluated for its suitability to help determine cytoarchitectonic boundaries by documenting the condition of each tissue section on a spreadsheet. Sections deemed suitable for use were photographed as described in *Section 2.6.1*. Team members (GPT, JSP) provisionally assigned an atlas level to the section by closely examining and documenting the nearest-neighbor spatial relations of identifiable cell groups in the *R series* photomicrographs and compared these with regions within the *Brain Maps 4.0* atlas. The criteria used to assign a given region of interest (ROI) in a tissue section to a specific atlas level is detailed below (*Sections 2.7.2.1–2.7.2.3*). In situations where two-closely spaced sections corresponded to the same atlas level, only one of the sections was deemed the best to represent distribution patterns for that atlas level (*i.e*., only one section and its labeling patterns were used to map patterns to that particular level).

Once initial assignments were made to all *R series* photomicrographs, they were migrated into Adobe Illustrator (AI) 2022 and annotated manually. A transparent data layer overlay was created for each photomicrograph and boundaries were drawn around relevant cell groupings identifiable in the Nissl-stained patterns. The left and right sides of the sections were also carefully noted to allow for accurate registration of Nissl-defined boundaries with tissue series processed for immunofluorescence. This was achieved by recording asymmetry in the shapes of the tissue hemispheres, especially in certain key areas (*e.g*., area between the pyramidal tracts; cross-sectional profiles of ventrally-located blood vessels investing the tissue). In the progress of performing these annotations and parcellations, team members revisited the tabulation of the original atlas assignments and, when deemed necessary, revised them on the basis of their improved familiarity with the spatial characteristics of the tissue. This iterative process, which is crucial to allow ample time for and which is necessary to ensure both accuracy and precision, is noted here as a critical part of this workflow. For instances where a Nissl-stained section was missing or damaged and could not be used to gather information to help with plane-of-section analysis and atlas level assignments, the chemoarchitecture of the fluorescently-labeled tissue series and its associated fiducials were used instead. In some cases, visualization of the fluorescently-labeled tissue section under dark-field illumination was sufficient to provide team members with important fiducials to make the determinations needed. The specific criteria and considerations used to determine correct spatial context for each ROI is detailed in the *Supplemental Methods*.

#### 2.7.3 Standardized mapping

Images of immunolabeled tissue sections were imported into Adobe Illustrator (AI) and aligned precisely with the data layer containing the photomicrograph of the corresponding adjacent *R series* Nissl-stained tissue section that served as a cytoarchitectural reference (*described in Sections 2.4.1 and 2.5*). Region boundaries that had been drawn over the Nissl-photograph data layer were then superimposed on the underlying fluorescence image, enabling each image set to be registered to the proper level of the *Brain Maps 4.0 (BM4.0)* open-access atlas [71].

To plot immunolabeled elements onto *BM4.0* atlas templates in AI, fluorescently-labeled fiber tracts and perikaryon profiles were traced and marked with circles and line segments, respectively on a separate data layer. If these representations shared a common region boundary (*i.e*., if they were all within the same region), they were grouped together in their layer. This layer was then copied and pasted as a collective set of drawn elements into an underlying data layer containing the corresponding atlas level template with the corresponding brain region. This import of drawn elements was achieved by aligning the grouped elements to be in as close of a registration as possible with the bounded regions on the map. Alignments generally consisted of rotating the drawn patterns to orient correctly with the map’s regional boundaries. In this manner, maps of the DBH-, ChAT-, and phospho-ERK1/2-immunoreactive signals were created for each subject.

#### 2.7.4 Region-delimited quantitation and assessment of signal co-localization

##### 2.7.4.1 Counting procedures

The rationale for our counting procedures is provided for interested readers in our *Supplemental Methods*. A few points regarding our counting procedures are noted here. First, in keeping with recommendations furnished. Immunolabeled perikaryon profiles were manually counted and annotated in a manner similar to that reported previously by one of us [66], but digital transparent overlays were utilized rather than acetate sheets affixed to printed hard copies of the images. Specifically, images of sections with ROIs corresponding to *BM4.0* atlas levels 51, 67, and 69 were opened in Adobe Illustrator (AI) 2022 and arranged into files containing the superimposed merged-channel and single-channel images, each in a separate layer. Two additional layers were created for the manual annotation of DBH- and pERK1/2-labeled perikaryon profiles.

To mark the locations of DBH-labeled perikaryon profiles, the reviewer made visible the layer containing the single-channel image corresponding to DBH immunofluorescence, and used the *Ellipse Tool* to place white-colored ellipses over labeled perikaryon profiles in the DBH layer. Keeping the layer containing DBH perikaryon profile markings visible, the team member toggled the single-channel pERK1/2 image on and off, determining if each DBH+ perikaryon profile was single or double-labeled based on the profile’s visibility in one or both channels, respectively.

To record the double-labeled perikaryon profiles, the *Smart Guides* “snap-to-center” function in AI was used to place a black ellipse on exactly the centroid coordinate of the DBH+ profile marking on a ‘pERK1/2 + DBH’ sublayer of the ‘pERK1/2’ layer. When all co-labeled perikaryon profiles were identified in this manner, the pERK1/2 single-channel image was used to record profiles displaying only signal for pERK1/2 in a sublayer called ‘pERK1/2 only’. For annotations of images corresponding to levels 67 and 69 of *BM4.0*, the ChAT immunolabel readily identified the cholinergic motor neurons of the DMX; any perikaryon profiles that were DBH+ or pERK1/2+ in the vicinity of the DMX but not ChAT+ were not considered DMX neurons.

To obtain the perikaryon profile counts, the layer or sublayer was selected and the number of objects within that layer was recorded. The number of pERK1/2 + DBH perikaryon profiles and pERK1/2-only profiles were added to determine the total number of pERK1/2-labeled profiles, and the number of pERK1/2 + DBH perikaryon profiles was subtracted from the total number of DBH+ profiles to determine the number of DBH-only profiles.

##### 2.7.4.3 Correction of oversampling error

We adapted the method of Abercrombie [100] to correct for overestimation of our perikaryon profile counts as detailed in the *Supplemental Methods*.

#### 2.7.5 Statistical analysis

The pERK1/2^+^, DBH^+^ and ChAT^+^ bilateral perikaryon profiles within the LC, NTSm and DMX were tallied on the sections corresponding to *BM4.0* atlas levels 51, 67, and 69. The average of the bilateral perikaryon profile counts [(left side counts + right side counts)/2] for each region, from each of the saline- and 2-DG-treated rats, was considered as a representative number of immunoreactive cells. The number of cells from each brain was used as a dependent variable, with saline or 2-DG treatment as the independent variable for statistical and exploratory data analyses. A Shapiro-Wilk test [101] was performed to determine the Gaussian distribution of the dependent variable in each treatment group. Phospho-ERK1/2^+^ perikaryon profile counts in the NTSm at levels 67 and 69 in 2-DG-treated rats did not meet the Gaussian distribution assumption at a *p*-value of 0.05. Since the distributions were not normal, and to be consistent throughout the study, non-parametric statistical tests were used to analyze all differences in profile counts across the sampled brain regions in this study.

Significance of the difference in the distributions of immunoreactive perikaryon profile counts between saline- and 2-DG-treated rats was analyzed for each region using the Wilcoxon Rank Sum test, with two-sided significance [102, 103]. A *p*-value less than or equal to 0.05 was considered statistically significant. False discovery rate was controlled by calculating Benjamini-Hochberg [104] adjusted *p*-values, with an FDR of 5% or below considered as statistically significant. Effect sizes were calculated and used to interpret the magnitude of the difference between saline- and 2-DG-treated subjects. Effect sizes of 0.1 to less than 0.3, 0.3 to less than 0.5, and greater than or equal to 0.5 were categorized as small, moderate, and large magnitudes, respectively. RStudio (R, version 3.6.2) [105] with *rstatix* [106] and *ggplot2* [107] packages were used for statistical analysis and to construct box plots and pie charts.

## 3. Results

The results of this study are divided into two main parts. First, rostral *rhombic brain (His, 1893)* [21] (RB) data for saline- and 2-DG-treated subjects are presented, specifically for the *locus ceruleus (Wenzel & Wenzel, 1912)* [83] (LC). Second, caudal RB data are considered, similarly comparing labeling patterns for the two groups for the *nucleus of solitary tract (>1840)* (NTS) and *dorsal motor nucleus of vagus nerve (>1840)* (DMX). For each part, individual subjects are first showcased to focus on paired comparisons of histological results, individual patterns of immunoreactivities in atlas maps, and morphological considerations of the signals. This is followed by group-level statistical comparisons of immunoreactive perikaryon profile counts between saline- and 2-DG-treated subjects and counts of activated perikaryon profiles sorted by cellular phenotype.

### 3.1. Cellular activation profiles in the LC for saline- and 2-DG-treated subjects

#### 3.1.1 Representative results in single pairs of subjects

##### 3.1.1.1 Phospho-ERK1/2-immunoreactive patterns mapped to Level 51

**Figure 2** shows the mapped distributions of phospho-ERK1/2-immunoreactive neurons in a portion of the LC that we sampled from subjects receiving intravenous saline or 2-DG injection. To identify the appropriate rostrocaudal locations of the tissue sections in both groups, we were guided by fiducial landmarks within the tissue, either visualized in the same section under dark-field illumination **(Fig. 2A-i)**, or within a thionine-stained section from an adjacent series of tissue collected from the same subject (*i.e*., *R series*; *see Methods*) **(Fig. 2B-i)**. The reference tissue allowed us to map the locations of the profiles observed in the fluorescence images to Level 51 of the *BM4.0* atlas [82]. Saline treatment was not associated with an appreciable number of phospho-ERK1/2^+^ perikaryon profiles in the LC **(Fig. 2A-iii)**. In contrast, a robust elevation in the numbers of phospho-ERK1/2^+^ perikaryon profiles was observed following 2-DG treatment **(Fig. 2B-iii)**. Virtually all of the phospho-ERK1/2 immunoreactivity appeared to be within DBH-immunoreactive neurons **(Fig. 2B-iv)**, a cellular phenotype well-documented to be abundant in this structure (*e.g*., [108–113]); these double-labeled perikaryon profiles were mapped as *gold-colored* circle glyphs on the corresponding atlas map **(Fig. 2B-v)**. At this level for this subject, all phospho-ERK1/2^+^ profiles appeared to be double-labeled for DBH-immunoreactivity, since none appeared to contain only the single label for phospho-ERK1/2 alone.

**Figure 2.**
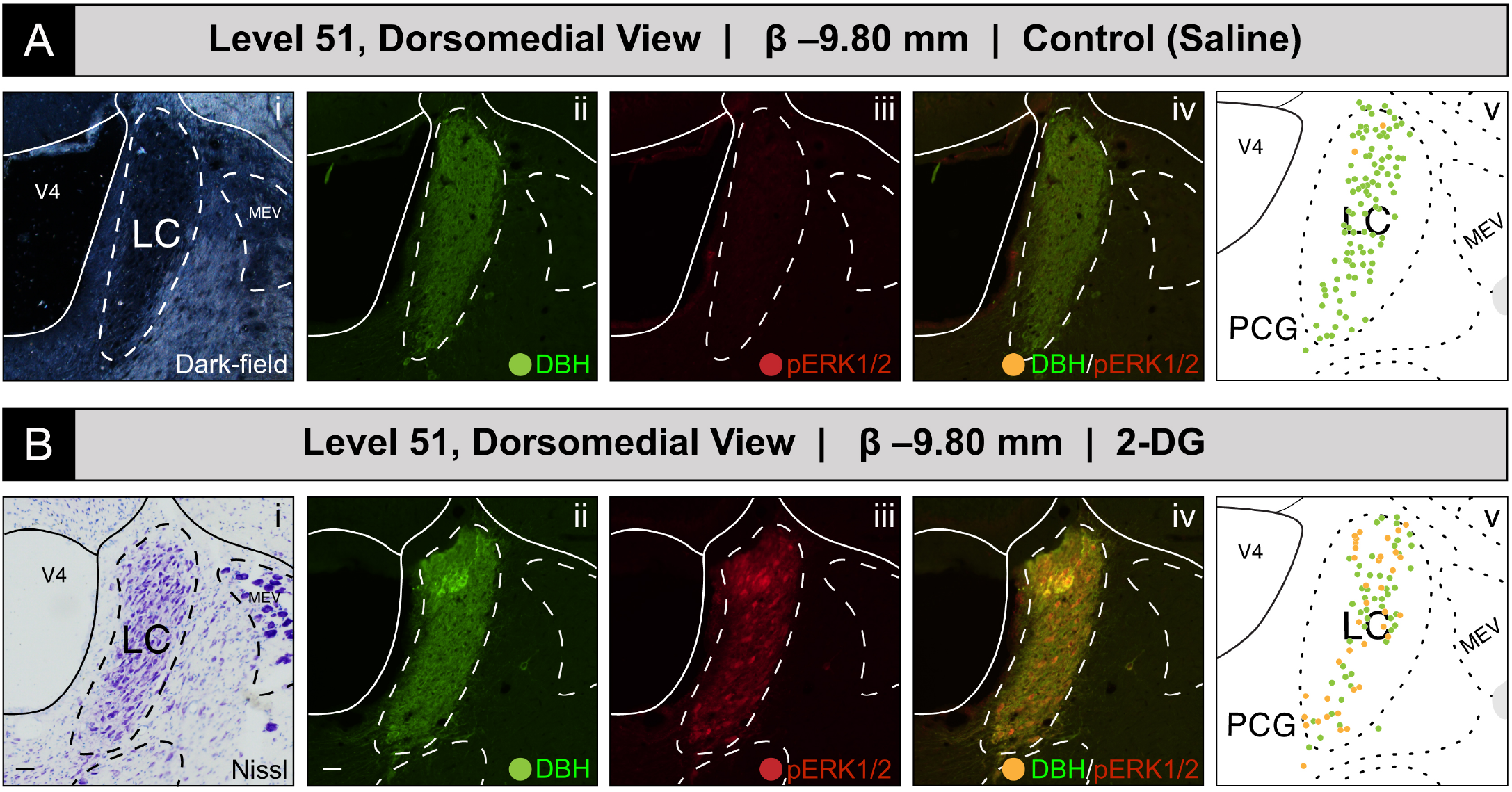
Increased recruitment of *locus ceruleus (Wenzel & Wenzel, 1912)* [83] (LC) neurons 15 min after intravenous 2-deoxy-D-glucose (2-DG) administration, as deduced from a comparison of LC perikaryon profiles in coronal-plane tissue sections from saline-**(A)** and 2-DG-treated **(B)** subjects. **(A, i–iv)**: Subject K10-012 (saline-treated). The LC of the right hemisphere, as visible in a dark-field image **(A-i)**, was immunostained for DBH (*green*; **A-ii**) and for the activation marker, phospho-ERK1/2 (*red*; **A-iii**), but displayed virtually no signal for the latter molecule. **(B, i–iv)**: Subject K10-026 (2-DG-treated). In contrast, this subject displayed robust phospho-ERK1/2-immunoreactive signal in the LC **(B-iii)**, the boundaries of which were determined using a Nissl-stained tissue section from an adjacent tissue series collected from the same subject (bright-field photomicrograph shown in **B-i**). The maps in **A(v)** and **B(v)** are maps of the LC from the *BM4.0* reference atlas [82]. Scale bars in **A-i/B-i** and **A-ii/B-ii** each mark 20 μm, and apply to the rest of the photomicrographs shown in this figure. Note for **A-i** that the dark-field image of the same tissue reacted with immunofluorescence was used to identify boundaries for the LC in the brain of subject K10-012, rather than an adjacent Nissl-stained section, which was damaged during tissue processing.

##### 3.1.1.2 Phospho-ERK1/2+ and DBH+ perikaryon profiles at other levels of the LC

Although we did not formally chart patterns of immunoreactivity to *BM4.0* atlas templates for tissue sections containing the LC at levels other than Level 51, we did observe phospho-ERK1/2^+^ profiles throughout the LC at all rostrocaudal levels containing this structure, primarily within 2-DG-treated subjects. For example, for another 2-DG-treated subject (#K10-027), perikaryon profiles in the LC again showed robust immunoreactivity for the activation marker in tissue sections just caudal to that mapped at Level 51 **(Figure 3)**. Specifically, at a rostrocaudal location that maps ~480 μm further caudally to a tissue plane that falls between *BM4.0* atlas levels 52 and 53, phospho-ERK1/2^+^ perikaryon profiles were again evident, especially in those situated dorsally within the structure **(Fig. 3B)**. Viewed under the higher power of a ×20 magnification objective lens and captured as a single-plane confocal scan, the DBH-immunoreactive signal could be observed within perikarya and neurite extensions, as has been reported by others [108–113]. Some of these neurites were oriented in parallel to the lateral margin of the *fourth ventricle principal part (Swanson, 2018)* (V4) **(Fig. 3A)**. Although double-labeled perikaryon profiles are evident at this level **(Fig. 3D)**, the relative abundance, brightness and saturation of the Alexa-488 fluorescence swamped the dimmer Cy3 signal of the phospho-ERK1/2 immunoreactivity in these profiles. These patterns were more readily observable if the opacity of the Alexa-488 signal was set to 35% without altering the Cy3 signal (**Fig. 3C**; compare profiles marked by arrows in panels **C** and **D**). In the double-labeled neurons, the presence of phospho-ERK1/2 immunoreactivity in the nuclear compartment of some perikarya was clearly evident, with a notable absence of fluorescent signal marking the presence of the nucleolus in many of these profiles **(Fig. 3D)**.

**Figure 3.**
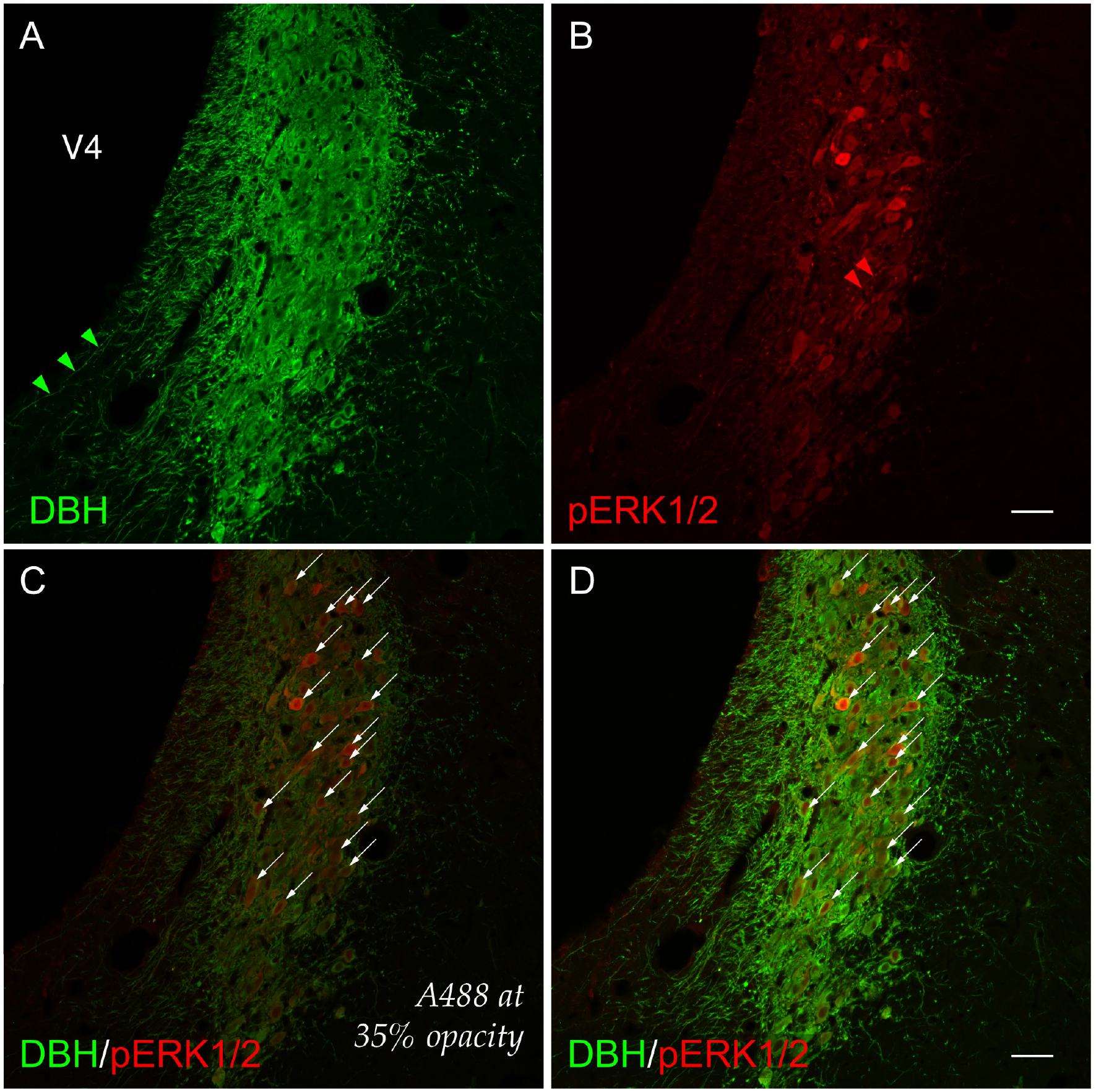
A higher-magnification view (cf. Fig. 2) of cellular activation observed in the *locus ceruleus (Wenzel & Wenzel, 1812)* (LC) [83] 15 min after 2-DG administration. Subject K10-027. **(A)** LC perikaryon profiles labeled for dopamine β-hydroxylase (DBH, *green*), and **(B)** phospho-ERK1/2 (pERK1/2, *red*) appear to co-label the same perikaryon profiles (**D**, *arrows*). **(C)** Co-labeled perikaryon profiles in the LC (identically marked by *arrows* as in **D**) with the A488 (*green*) channel set at 35% opacity renders visible the subtle, graded activation profiles of phospho-ERK1/2^+^ signal. *Arrowheads* (*green* in **A**; *red* in **B**) mark single-labeled neurites immunoreactive for DBH or pERK1/2, respectively. Scale bars in **B and D** mark 50 μm and apply to all photomicrographs.

#### 3.1.2 Group-level effects of saline versus 2-deoxy-D-glucose administration on LC activation

##### 3.1.2.1 Quantitative analysis of LC activation

To determine LC activation in association with saline and 2-DG treatments, pERK1/2- and DBH-immunoreactive perikaryon profiles were counted in the LC at *BM4.0* atlas level 51 for all subjects. **Figure 4** shows box plots of Abercrombie-corrected phospho-ERK1/2^+^ and DBH^+^ perikaryon profile counts and **Supplemental Table 1 (Table S1)** presents the descriptive statistics of our quantitative analysis. Median numbers of phospho-ERK1/2-immunoreactive profiles in saline and 2-DG groups were 15 and 33, respectively **(Table S1)**, and the distributions in the two groups differed significantly (Wilcoxon statistic = 25, n_1_ = n_2_ = 5, FDR-adjusted *p* < 0.05; **Table S2**). Specifically, the number of pERK1/2-immunoreactive profiles displayed in the LC was significantly elevated for 2-DG-compared to saline-treated rats (**Fig. 4**, *‘total’ column* (pERK1/2: +); FDR-adjusted *p* < 0.05; effect size: large; **Table S2**). Similarly, median numbers of double-labeled profiles in saline and 2-DG groups were 13 and 33, respectively **(Table S1)**, with distributions in the two groups that again were significantly different (Wilcoxon statistic = 25, n_1_ = n_2_ = 5, FDR-adjusted *p* < 0.05; **Table S2**). Thus, the double-labeled counts of LC profiles of 2-DG rats were elevated significantly compared to those of saline-treated rats (**Fig. 4**, *‘double-labeled’ column* (pERK1/2, DBH: both +); effect size: large; **Table S2**). In contrast, there were no statistically significant differences in DBH perikaryon profile counts between saline- and 2-DG-treated rats (**Fig. 4**, *‘total’ column* (DBH: +) and *‘single-labeled’ column* (DBH: +, pERK1/2: –); see also **Table S2**).

**Figure 4.**
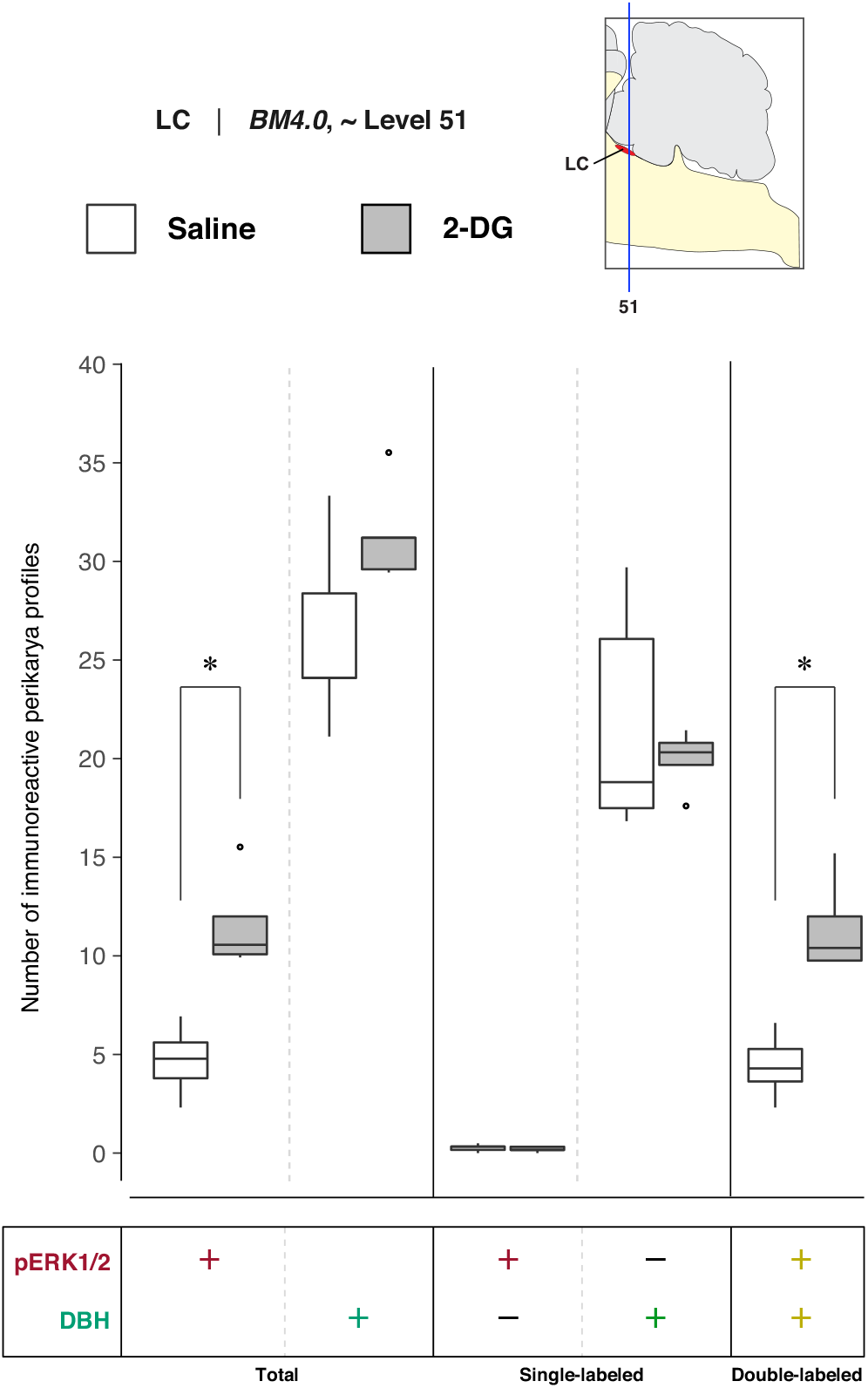
Increased recruitment of *locus ceruleus (Wenzel & Wenzel, 1812)* (LC) [83] neurons 15 min after 2-DG administration. Abercrombie-corrected [100] perikaryon profile counts from both saline- and 2-DG-treated subjects are represented as box plots displaying the total numbers of phospho-ERK1/2- or DBH-immunoreactive profiles, or only the single- and double-labeled profiles, from tissue sections identified to be in register with *BM4.0* atlas level 51 (map, *upper right*). An asterisk (*) denotes statistical significance between groups, FDR-adjusted *p* < 0.05. See **Tables S1 & S2** for details.

##### 3.1.2.2 Relative differences in the numbers of DBH-immunoreactive perikaryon profiles in the LC of saline- and 2-DG-treated subjects

To determine whether 2-DG treatment was associated with changes in the numbers of DBH-immunoreactive neurons in the LC, the relative percentages of Abercrombie-corrected single (pERK1/2^+^ only or DBH^+^ only) and double-labeled (pERK1/2^+^ & DBH^+^) perikaryon profiles were calculated for saline and 2-DG groups, and visualized as pie charts in **(Supplemental) Figure S1**. Consistent with our microscopic observations, a majority of the pERK1/2+ perikaryon profiles in LC were identified to be DBH+ in both saline-(94.9%) and 2-DG-treated (98.3%) rats (**Fig. S1**, pERK+ row, pERK1/2^+^ & DBH^+^; *yellow-colored fraction*). For the total numbers of DBH^+^ perikaryon profiles, a higher percentage of pERK1/2^+^ profiles were observed in the LC of 2-DG-(36.4%) than saline-treated (16.9%) rats (**Fig. S1**, DBH+ row, pERK1/2^+^ & DBH^+^, *yellow-colored fraction*). In contrast, the average total numbers of DBH^+^ perikaryon profiles were not significantly different between the two groups (FDR-adjusted *p*=0.22; **Table S2**).

### 3.2. Cellular activation profiles in the NTS and DMX for saline- and 2-DG-treated subjects

#### 3.2.1 Representative results in single pairs of subjects at atlas level 67

##### 3.2.1.1 Phospho-ERK1/2-immunoreactive patterns mapped to Level 67

**Figure 5** shows the mapped distributions of phospho-ERK1/2-immunoreactive signal in activated NTS and DMX neurons following saline or 2-DG administration. Saline treatment was not associated with a substantial amount of phospho-ERK1/2-immunopositive perikaryon profiles in the NTS at Level 67 **(Fig. 5A-iii)**. In comparison, 2-DG treatment was associated with a marked increase in phospho-ERK1/2 immunoreactive perikaryon profiles within the NTSm **(Fig. 5B-iii)**. This elevation correlated to both single-labeled phospho-ERK1/2 profiles and those double-labeled for DBH-immunoreactivity **(Fig. 5B-vi)**, that were represented as *red-* and *yellow-*colored circle glyphs, respectively, on the corresponding atlas map **(Fig. 5B-v)**.

**Figure 5.**
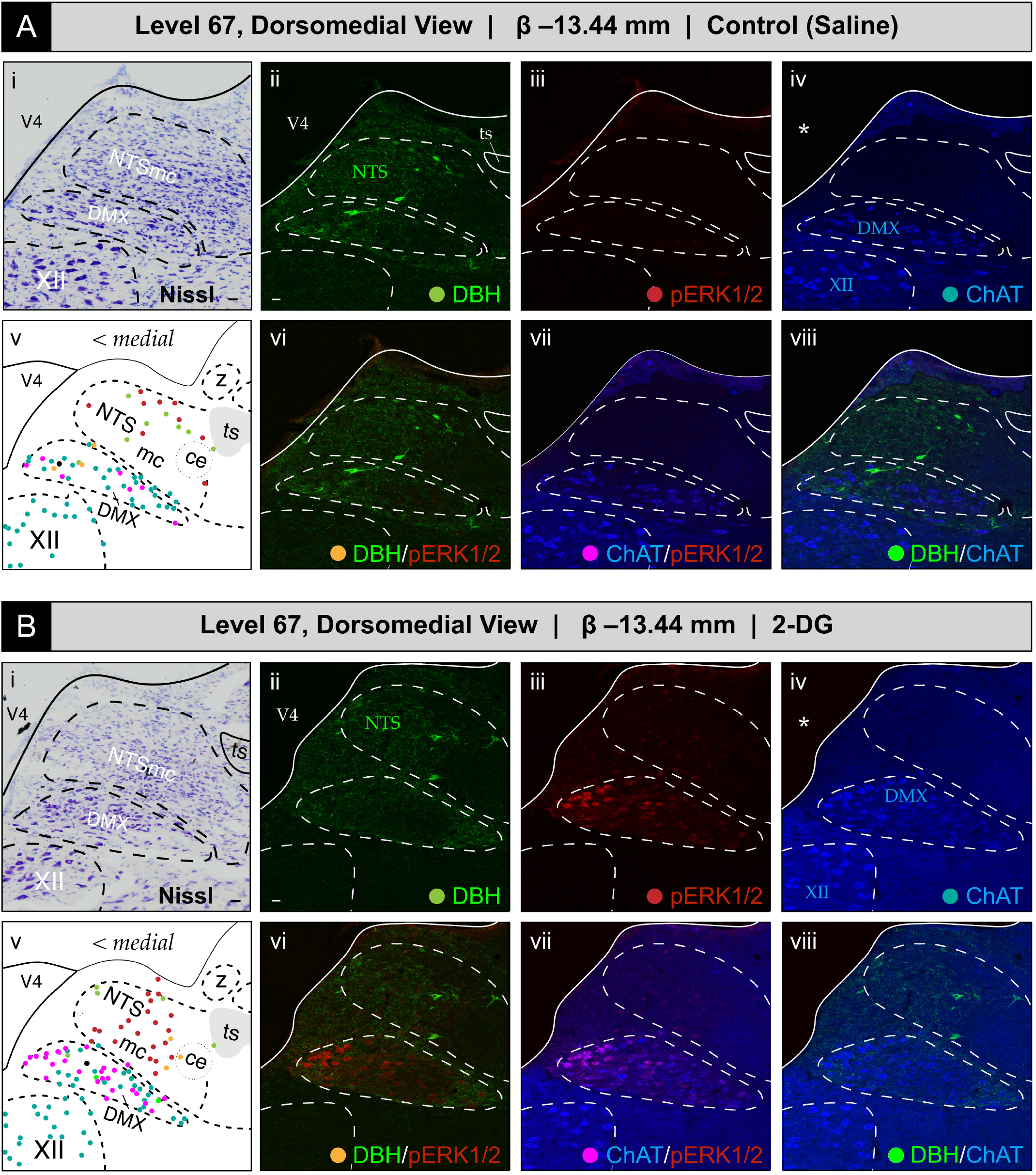
Glycemic challenge is associated with the recruitment of dorsal *medulla (Winslow, 1733)* [37] (MY) neurons mapped to atlas level 67 of *Brain Maps 4.0 (BM4.0)* [82]. Cytoarchitecture- and chemoarchitecture-based spatial analysis of distributed phosphorylated MAP kinase (pERK1/2) signal within dorsal RB neurons at the level of the *nucleus of solitary tract (>1840)* (NTS) (Level 67 in the *BM4.0* rat brain atlas). **(A, B)** The panels display bright-field Nissl photomicrographs **(i)**, single-**(ii–iv)** and dual-channel **(vi–viii)** fluorescence images, and *BM4.0* atlas maps **(v)** for saline-**(A)** and 2-DG-treated **(B)** subjects of individual immunoreactivity patterns for dopamine β-hydroxylase (DBH, *green*), phospho-ERK1/2 (pERK1/2, *red*) and choline acetyltransferase (ChAT, *blue*). The patterns fall near and within the boundaries for the NTS, *dorsal motor nucleus of vagus nerve (>1840)* (DMX), and *hypoglossal nucleus (>1840)* (XII). Note that the NTS label is colored *green* to signify its boundary determination using the pattern of immunoreactivity observed for DBH in the green channel, and DMX and XII labels are colored *blue* to signify their boundary determinations using ChAT-immunoreactive signal in the far-red channel. Tissue is shown in the coronal plane. Scale bars in **(i)**and **(ii)**, which mark 20 μm, and the medial orientation noted for the maps in **(v)**, apply to all photomicrographs. *Asterisks* in **A-iv** and **B-iv** mark the locations of masks placed over the ventricular space using Adobe Photoshop, to cover visually-distracting autofluorescence observed under the far-red channel from the mounting medium; this mask is used for all images containing far-red (*blue-colored*) signal (thus, this also applies to **A-vii/viii** and **B-vii/viii**). Headers for **A** and **B** note the inferred anteroposterior stereotaxic coordinate, expressed as millimeters from the Bregma (β) skull suture. See the list of abbreviations at the end of this article for a description of those marked on the maps.

ChAT-immunopositive neurons in the DMX, depicted in *blue*, displayed low levels of basal activation in saline-treated subjects **(Fig. 5A-vii)**, and appear as *magenta* circles on the atlas maps when also immunoreactive for phospho-ERK1/2. In some subjects, the numbers of phospho-ERK1/2^+^ perikaryon profiles in the DMX were elevated following 2-DG but not saline treatment **(Fig. 5B-vii)**, but as noted in *Section 3.2.2(c)*, this difference did not reach statistical significance at a group level. In some instances, ChAT-labeled perikaryon profiles were observed to also be DBH-immunopositive, shown as *neon green* circles, and represented as *black* circles when immunoreactive for all three markers. In contrast, no phospho-ERK1/2 immunoreactivity was observed in the *hypoglossal nucleus (>1840)* (XII), and few or no DBH-immunoreactive fibers were observed within the *nucleus of solitary tract, lateral part (>1840)*.

#### 3.2.2 Group-level effects of saline versus 2-deoxy-D-glucose administration on NTS and DMX activation at BM4.0 Atlas level 67

##### 3.2.2(a) Quantitative analysis of NTS activation at level 67

To determine NTS activation in association with saline and 2-DG treatments, pERK1/2- and DBH-immunoreactive perikaryon profiles were counted in the NTS at *BM4.0* atlas level 67 for all subjects (n = 6 for the saline-treated group; n = 5 for the 2-DG-treated group). **Figure 6** shows box plots of Abercrombie-corrected phospho-ERK1/2^+^ and DBH^+^ perikaryon profile counts. **Tables S3 and S4** present the descriptive and test statistics, respectively, of our quantitative analysis.

**Figure 6.**
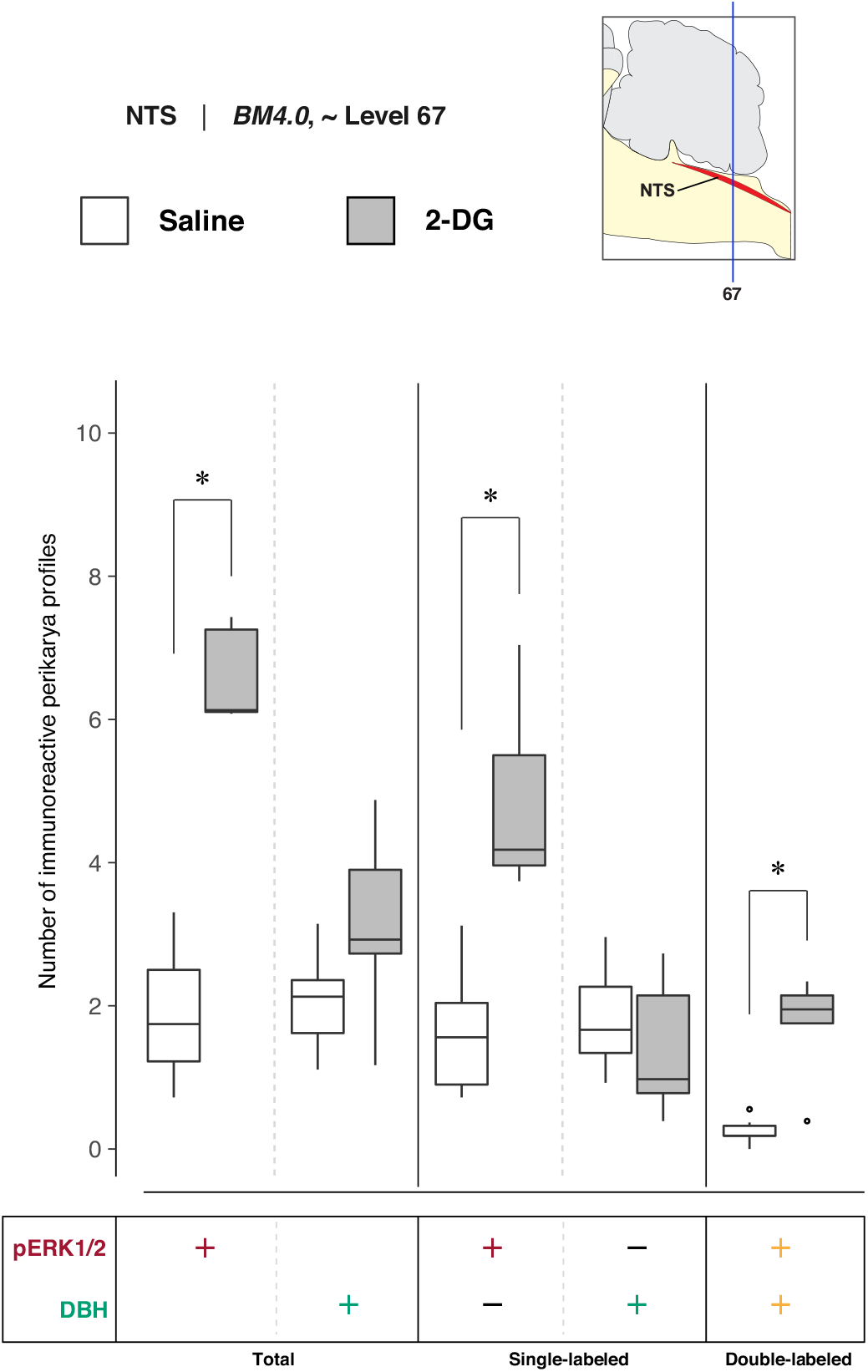
Increased recruitment of *BM4.0* atlas level 67-registered *nucleus of solitary tract (>1840)* (NTS) neurons 15 min after 2-DG administration. Abercrombie-corrected [100] perikaryon profile counts from both saline- and 2-DG-treated subjects are represented as box plots displaying the total numbers of phospho-ERK1/2- or DBH-immunoreactive profiles, or only the single- and double-labeled profiles, from tissue sections identified to be in register with *BM4.0* atlas level 67 (map, *upper right*). An asterisk (*) denotes statistical significance between groups, FDR-adjusted *p* < 0.05. See **Tables S3 & S4** for details.

Median numbers of pERK1/2-immunoreactive profiles in saline and 2-DG groups were 1.7 and 6.1, respectively **(Table S3)**, and the distributions in the two groups differed significantly (Wilcoxon statistic = 30, FDR-adjusted *p* < 0.05; **Table S4**). The number of pERK1/2-immunoreactive profiles within the NTS significantly increased following 2-DG treatment as compared to saline-treated controls (**Fig. 6**, *‘total’ column* (pERK1/2: +); FDR-adjusted *p* < 0.05; effect size: large; **Table S4**). The median numbers of double-labeled profiles in saline and 2-DG groups were 0.2 and 1.9, respectively **(Table S3)**, that maintained significantly different distributions between groups (Wilcoxon statistic = 29, FDR-adjusted *p* < 0.05; **Table S4**). The elevated counts observed in 2-DG-treated rats translated across single-(**Fig. 6**, *‘single-labeled’ column* (pERK1/2: +, DBH: –); FDR-adjusted *p* < 0.05; effect size: large; **Table S4**) and dual-labeled perikaryon profile counts (**Fig. 6**, *‘double-labeled’ column* (pERK1/2: +, DBH: +); FDR-adjusted *p* < 0.05; effect size: large; **Table S6**). Saline- and 2-DG-treated rats did not significantly differ in the total DBH-immunoreactive profile counts (**Fig. 6**, *‘total’ column* (DBH: +); see also **Table S4**).

##### 3.2.2(b) Relative differences in the numbers of DBH-immunoreactive perikarya in the NTS at level 67 of saline- and 2-DG-treated subjects

To determine whether 2-DG treatment was associated with changes in the numbers of DBH-immunoreactive neurons in the NTS at level 67, the relative percentages of Abercrombie-corrected single- (pERK1/2^+^ only or DBH^+^ only) and double-labeled (pERK1/2^+^ & DBH^+^) perikaryon profiles were calculated for saline and 2-DG groups and visualized as pie charts in **Figure S2**. In contrast to relative percentages observed in the LC at atlas level 51, the majority of the pERK1/2^+^ perikaryon profiles in the NTS at level 67 were identified to be DBH^−^ in both saline- (86.9%) and 2-DG-treated (74%) rats (**Fig. S2**, pERK+ row, pERK1/2^+^ only; *maroon-colored fraction*). For total numbers of DBH+ perikaryon profiles, a higher percentage of pERK1/2^+^ profiles were observed in the NTS of 2-DG- (55%) than saline-treated (11.9%) rats (**Fig. S2**, DBH+ row, pERK1/2^+^ & DBH^+^; *yellow-colored fraction*).

##### 3.2.2(c) Quantitative analysis of DMX activation at level 67 and relative differences in the numbers of DBH- and ChAT-immunoreactive perikarya

To determine DMX activation in association with saline and 2-DG treatments, pERK1/2-, ChAT-, and DBH-immunoreactive perikaryon profiles were counted in the DMX at *BM4.0* atlas level 67 for all but one subject (n = 6 for the saline-treated group, n = 5 for the 2-DG-treated group; tissue for the 2-DG-treated subject, K10-008, was not available at this level). **Figure 7** shows box plots of Abercrombie-corrected phospho-ERK1/2^+^, ChAT^+^, and DBH^+^ perikaryon profile counts for these eleven subjects. **Tables S3 and S4** present the descriptive statistics and test statistics, respectively, of all comparisons at this level for these subjects. Although we observed treatment-associated differences in Abercrombie-corrected profiles displaying pERK1/2 immunoreactivity in the DMX at *BM4.0* atlas level 67 in some subjects **(Fig. 5)**, at a group level, the numbers of profiles (single-, double-, or triple-labeled) did not differ significantly between saline- and 2-DG-treated rats (see **Fig. 7** and **Table S4**). Similarly, no differences were observed in the numbers of ChAT^+^ and DBH^+^ perikaryon profiles between treatment groups.

**Figure 7.**
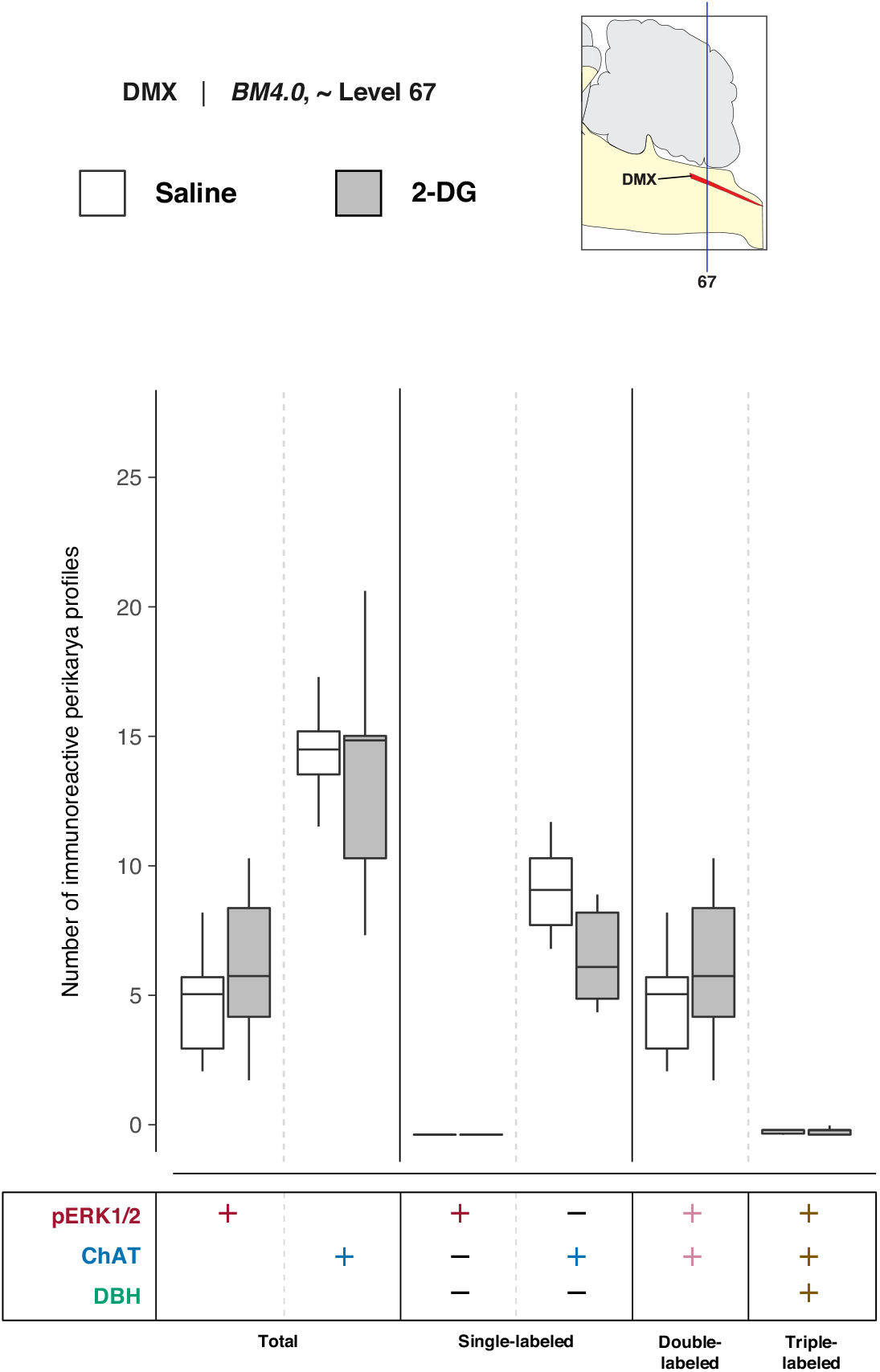
2-DG administration, at a group level, is not associated with an alteration of activation patterns in *dorsal motor nucleus of vagus nerve (>1840)* (DMX) perikaryon profiles registered to *BM4.0* atlas level 67. Abercrombie-corrected [100] perikaryon profile counts from both saline- and 2-DG-treated subjects are represented as box plots displaying the total numbers of phospho-ERK1/2- or ChAT- or DBH-immunoreactive profiles, or only the single-, double-, and triple-labeled profiles, from tissue sections identified to be in register with *BM4.0* atlas level 67 (map, *upper right*). Differences in the numbers of perikaryon profiles were not significant; see **Tables S3 & S4** for a statistical summary.

In the absence of significant changes in the number of DMX perikaryon profiles, the relative percentages of Abercrombie-corrected single-, double-, and triple-labeled profiles were calculated for saline and 2-DG groups, and visualized as pie charts in **Figure S3**. The majority of the pERK1/2^+^ perikaryon profiles in the DMX at level 67 were identified to be ChAT^+^ but negative for DBH were in the overwhelming majority of profiles in both saline- (97.7%) and 2-DG-treated (97.8%) rats (**Fig. S3**, pERK+ row, pERK1/2^+^ & ChAT^+^; *magenta-colored fraction*). Interestingly, for the total number of ChAT^+^ perikaryon profiles, the percentage of pERK1/2^+^ & ChAT^+^ double-labeled perikaryon profiles appears to be greater in 2-DG- (45%) than saline-treated (33.9%) rats.

#### 3.2.3 Representative results in single pairs of subjects at atlas level 69

##### 3.2.3.1 Phospho-ERK1/2-immunoreactive patterns mapped to Level 69

**Figure 8** shows the mapped distributions of phospho-ERK1/2 immunoreactivity in NTS and DMX perikaryon profiles at level 69 following saline or 2-DG administration. Saline treatment was associated with a relatively low number of phospho-ERK1/2-immunoreactive perikaryon profiles in the NTS at level 69 **(Fig. 8A-iii)**. Conversely, 2-DG treatment corresponded to pronounced numbers of immunoreactive profiles in comparable tissue sections **(Fig. 8B-iii)**. Saline treatment was also associated with a low amount of basally-activated ChAT-immunopositive perikarya that dimly emitted phospho-ERK1/2 signal in the DMX at level 69 **(Fig. 8A-vii)**. Similar to level 67, the total numbers of phospho-ERK1/2-immunoreactive perikaryon profiles, and relative brightness of the fluorescent signal, increased to a slight extent following 2-DG administration **(Fig. 8B-vii)**. In both groups, fewer ChAT-immunoreactive profiles that were also DBH-immunopositive, or triple-labeled, were observed at this level. A few phospho-ERK1/2-immunoreactive profiles were found in the NTSco, but not in the NTSl or XII.

**Figure 8.**
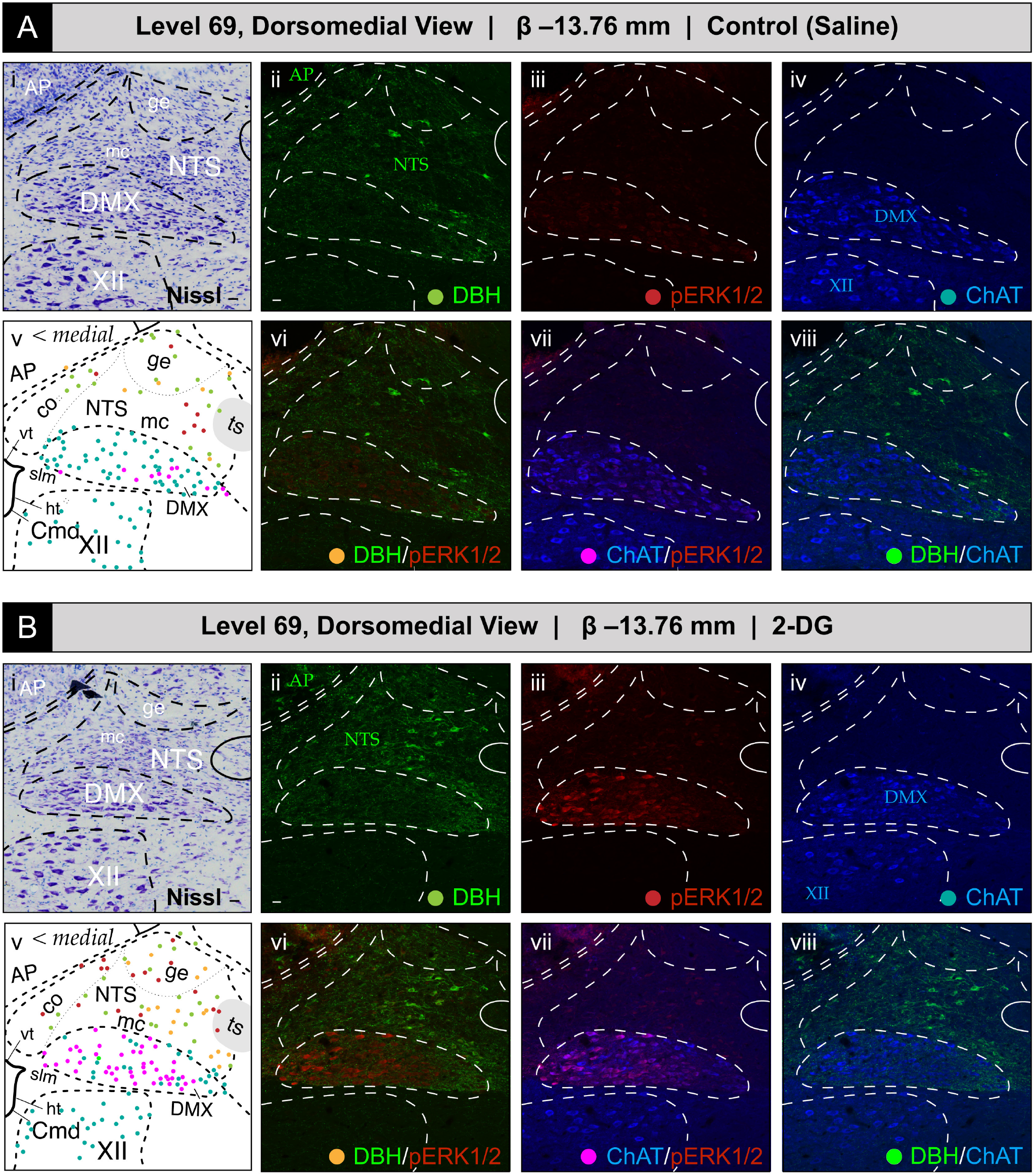
Glycemic challenge is associated with the recruitment of dorsal *medulla (Winslow, 1733)* [37] (MY) neurons mapped to atlas level 69 of *Brain Maps 4.0 (BM4.0)* at the level of the *area postrema (>1840)* (AP) [82]. Cytoarchitecture- and chemoarchitecture-based spatial analysis of immunoreactivity patterns for dopamine β-hydroxylase (DBH, *green*), phospho-ERK1/2 (pERK1/2, *red*) and choline acetyltransferase (ChAT, *blue*) within dorsal MY neurons. (**A, B**) The panels display coronal-plane Nissl photomicrographs (**i**), single- (**ii–iv**) and dual-channel (**vi–viii**) fluorescence images, and *BM4.0* atlas maps (**v**) for saline- (**A**) and 2-DG-treated (**B**) subjects. The patterns fall near or within the boundaries for the AP, *nucleus of solitary tract (>1840)* (NTS), *dorsal motor nucleus of vagus nerve (>1840)* (DMX), and *hypoglossal nucleus (>1840)* (XII). Colors for the AP, NTS, DMX, XII labels match those of the labeling patterns used to help define their boundaries. Scale bars in (**i**) and (**ii**), which mark 20 μm, and the medial orientation noted in (**v**), apply to all images. Headers list the inferred anteroposterior stereotaxic coordinate (in mm from the Bregma (β) skull suture). See the list of abbreviations at the end of this article for a description of those marked on the maps.

**Figure 9** presents confocal images of tissue sections that better showcase the elevated numbers of perikaryon profiles displaying phospho-ERK1/2 immunoreactivity within the NTS and DMX of 2-DG-treated subjects as compared to those receiving saline vehicle. Increased numbers of perikaryon profiles co-labeled with DBH immunoreactivity, and those expressing only phospho-ERK1/2 immunoreactivity, were observed in the NTS in association with 2-DG treatment **(Fig. 9B”-ii)**. Within the DMX, saline-treated subjects displayed few phospho-ERK1/2-immunoreactive profiles that mostly populated the medial portion of the nucleus **(Fig. 9A”-ii)**. 2-DG treatment was associated with minor elevations in phospho-ERK1/2-immunoreactive profiles within the DMX that did not appear to show a preferential distribution through the nucleus **(Fig. 9B”-ii)**.

**Figure 9.**
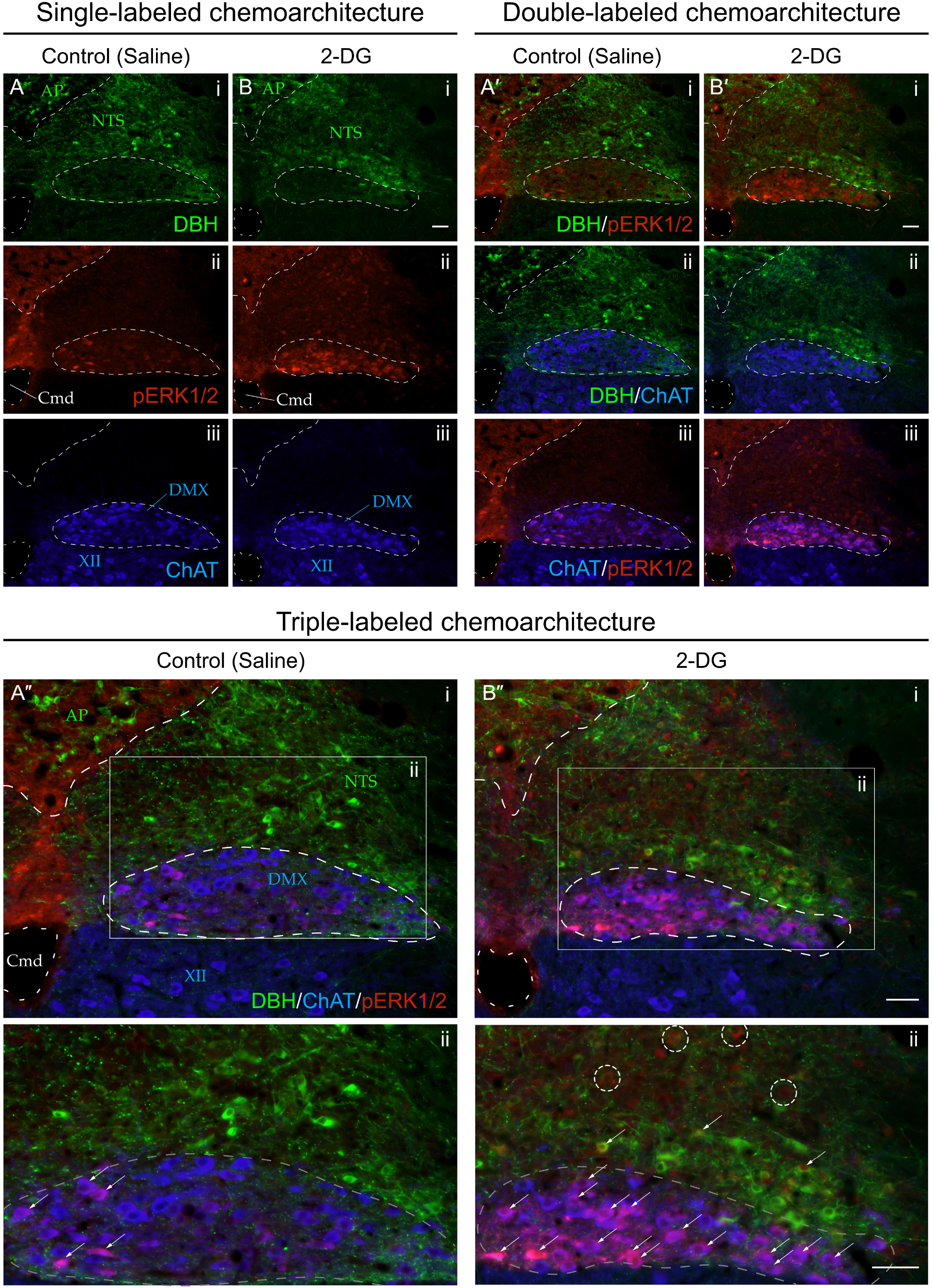
(*on previous page*). Increased recruitment of dorsal *medulla (Winslow, 1733)* [37] (MY) neurons 15 min after intravenous 2-deoxy-D-glucose (2-DG) administration relative to saline-treated subjects. Single-channel **(A, B)**, dual-channel **(A’, B’)**, and triple-channel **(A’’, B’’)** coronal-plane images, from saline- **(A, A’, A’’)** and 2-DG-treated **(B, B’, B’’)** subjects, of individual immunoreactivity patterns for dopamine β-hydroxylase (DBH; *green*), phospho-ERK1/2 (pERK1/2; *red*) and choline acetyltransferase (ChAT; *blue*) in the *area postrema (>1840)* (AP), *nucleus of solitary tract (>1840)* (NTS), *dorsal motor nucleus of vagus nerve (>1840)* (DMX), and *hypoglossal nucleus (>1840)* (XII), in the vicinity of the *central canal of medulla (>1840)* (Cmd). Note that the AP and NTS labels are colored *green* to signify their boundary determinations using the pattern of immunoreactivity observed for DBH in the green channel, and DMX and XII locations are labeled in *blue* to signify their boundary determinations using ChAT immunoreactive signal in the far-red channel. See the methods section for more details. Phospho-ERK1/2 immunoreactivity (*red*) is also present in NTS perikaryon profiles of unidentified phenotype (see *dashed circles* in **B’’.ii**) in 2-DG-treated but not saline-treated subjects.

#### 3.2.4 Group-level effects of saline versus 2-deoxy-D-glucose administration on NTS and DMX activation at BM4.0 atlas level 69

##### 3.2.4(a) Quantitative analysis of NTS activation at level 69

To determine NTS activation in association with saline and 2-DG treatments, phospho-ERK1/2- and DBH-immunoreactive perikaryon profiles were counted in the NTS at *BM4.0* atlas Level 69 for saline-(n=6) and 2-DG-treated (n=4) subjects. **Figure 10** shows box plots of Abercrombie-corrected phospho-ERK1/2^+^ and DBH^+^ perikaryon profile counts. **Tables S5 and S6** present the descriptive and test statistics, respectively, of our quantitative analysis. Median numbers of phospho-ERK1/2-immunoreactive profiles in saline and 2-DG groups were 3.6 and 8.3, respectively **(Table S5)**, and the distributions in the two groups differed significantly (Wilcoxon statistic = 24, FDR-adjusted *p* < 0.05; effect size: large; **Table S6**). The number of phospho-ERK1/2-immunoreactive profiles within the NTS significantly increased following 2-DG treatment as compared to saline-treated controls (**Fig. 10**; ‘total’ column (pERK1/2: +); FDR-adjusted *p* < 0.05; effect size: large; **Table S6**). The median numbers of double-labeled profiles in saline and 2-DG groups were 0.8 and 2.9, respectively **(Table S5)**, that again were significantly different (Wilcoxon statistic = 24, FDR-adjusted *p* < 0.05; effect size: large; **Table S6**). Total numbers of DBH-labeled profiles (**Fig. 10**, ‘total’ column (DBH: +); see also **Table S6**) did not significantly vary between treatment groups.

**Figure 10.**
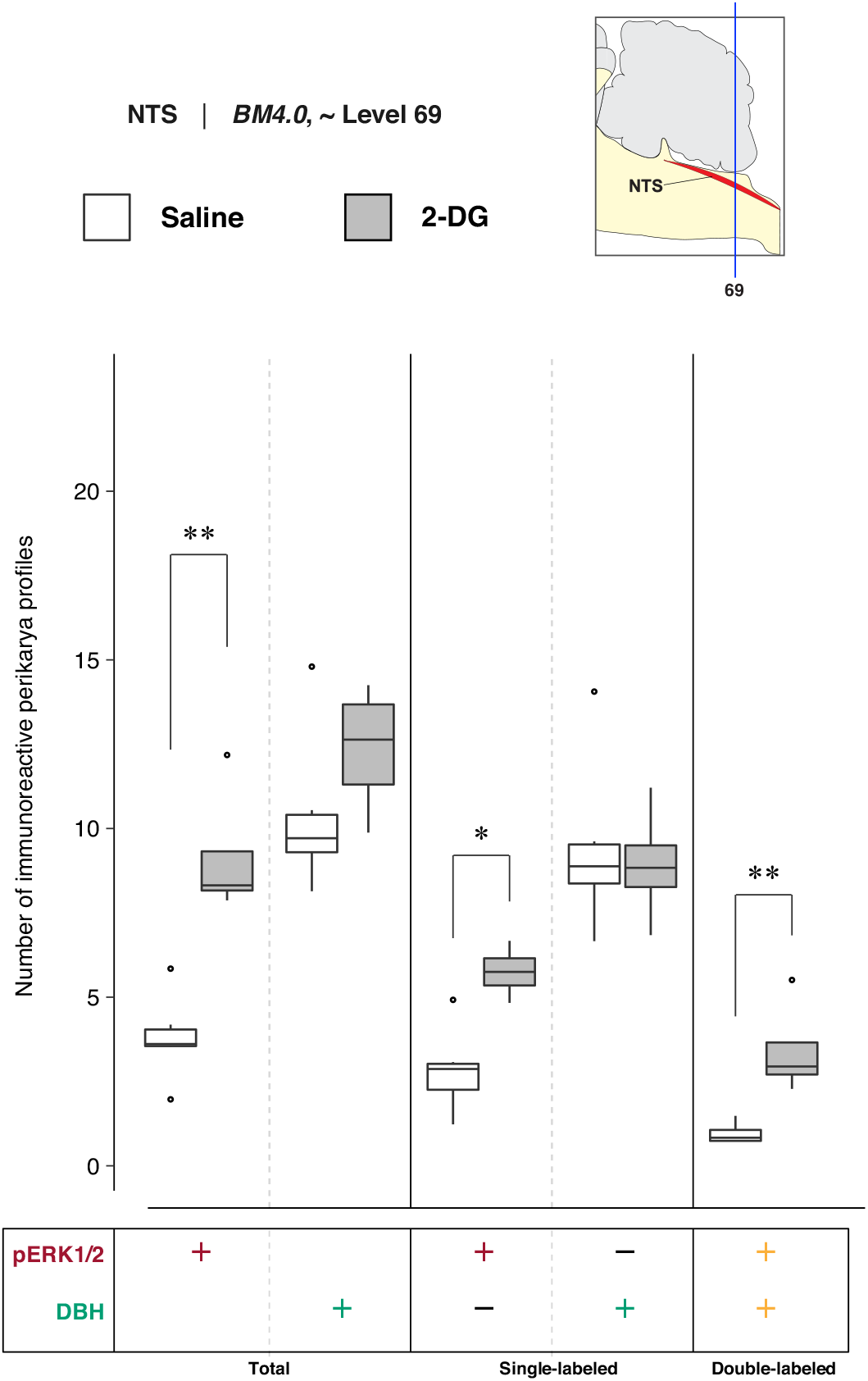
2-DG administration is associated with a marked alteration of activation patterns in *nucleus of solitary tract (>1840)* (NTS) perikaryon profiles registered to *BM4.0* atlas level 69. Abercrombie-corrected [100] perikaryon profile counts from both saline- and 2-DG-treated subjects are represented as box plots displaying the total numbers of phospho-ERK1/2- or DBH-immunoreactive profiles, or only the single- and double-labeled profiles, from tissue sections identified to be in register with *BM4.0* atlas level 69 (map, *upper right*). An asterisk (*) denotes statistical significance between groups with an FDR-adjusted *p* < 0.05; a double asterisk (**) denotes an FDR-sdjusted *p* < 0.01. See **Tables S5 & S6** for details.

##### 3.2.4(b) 2-DG-treatment associated relative changes in perikaryon phenotype composition of the NTS at level 69

To identify 2-DG treatment-associated changes in perikaryon phenotype composition in the NTS at level 69, the relative percentages of Abercrombie-corrected single (pERK1/2^+^ only or DBH^+^ only) and double-labeled (pERK1/2^+^ & DBH^+^) perikaryon profiles were calculated for saline and 2-DG groups, and visualized as pie charts in **Figure S4**. Similar to the relative percentages observed in the NTS region at atlas level 67, a majority of the pERK1/2^+^ perikaryon profiles were identified to be DBH^−^ in both saline- (74.8%) and 2-DG-treated (62.7%) rats (**Fig. S4**, pERK+ row, pERK1/2^+^ only; *maroon-colored* slice). Consistent with level 67 NTS perikaryon profile relative percentages, in DBH^+^ perikaryon profiles, a higher percentage of pERK1/2^+^ profiles was observed in the NTS region of 2-DG-(27.7%) than saline-treated (9.2%) rats (**Fig. S4**, DBH+ row, pERK1/2^+^ & DBH^+^; *yellow-colored* slice).

##### 3.2.4(c) Quantitative analysis of DMX activation and relative changes in perikaryon phenotype composition at level 69

To determine DMX activation in association with saline and 2-DG treatments, phospho-ERK1/2-, ChAT-, and DBH-immunoreactive perikaryon profiles were counted in the DMX at *BM4.0* atlas level 69 for saline-(n=6) and 2-DG-treated (n=4) subjects. **Figure 11** shows box plots of Abercrombie-corrected phospho-ERK1/2^+^, ChAT^+^, and DBH^+^ perikaryon profile counts. **Tables S5 & S6** present the descriptive and test statistics, respectively, of our quantitative analysis. Median numbers of phospho-ERK1/2-immunoreactive profiles in saline and 2-DG groups were 8.8 and 8.5, respectively **(Table S5)**, and the distributions in the two groups were not statistically different (Wilcoxon statistic = 11.5, FDR-adjusted *p* = 1; **Table S6**). The total numbers of ChAT-labeled profiles (**Fig. 11**, ‘total’ column (ChAT: +); see also **Table S6**) or ChAT-labeled profiles also immunopositive for DBH (**Fig. 11**; see also **Table S6**) did not vary significantly between treatment groups. Thus, triple-labeled perikaryon profile counts (**Fig. 11**, ‘triple-labeled’ column; see also **Table S6**) were not significantly different between groups.

**Figure 11.**
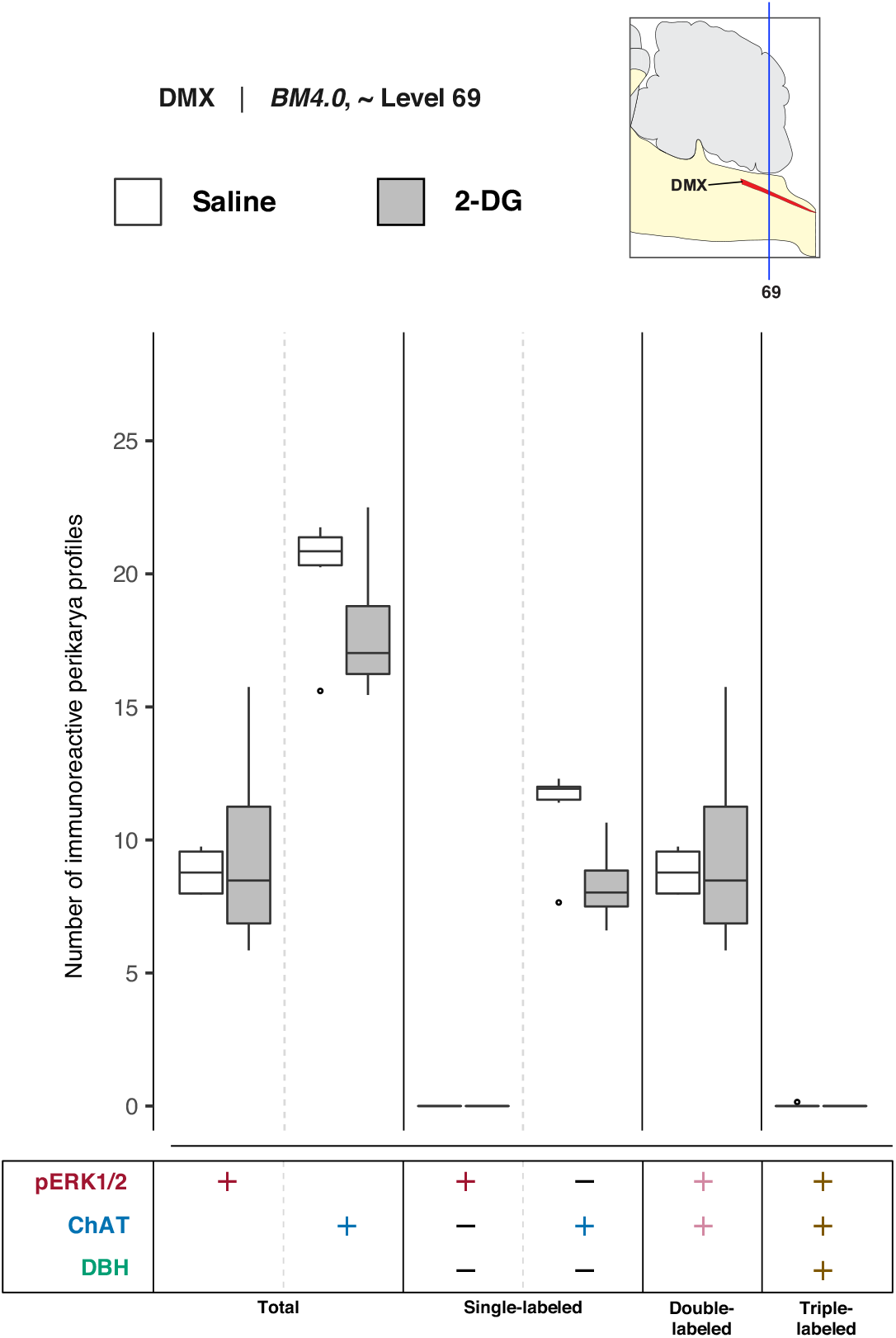
2-DG administration does not alter the activation patterns in *dorsal motor nucleus of vagus nerve (>1840)* (DMX) perikaryon profiles registered to *BM4.0* atlas level 69. Abercrombie-corrected [100] perikaryon profile counts from both saline- and 2-DG-treated subjects are represented as box plots displaying the total numbers of phospho-ERK1/2- or ChAT- or DBH-immunoreactive profiles, or only the single-, double-, and triple-labeled profiles, from tissue sections identified to be in register with *BM4.0* atlas level 69 (map, *upper right*). Perikaryon profile count differences were not significant; see **Tables S5 & S6** for a statistical summary.

In the absence of significant differences in the numbers of DMX perikaryon profiles, the relative percentages of Abercrombie-corrected single-, double-, and triple-labeled profiles were calculated for saline and 2-DG groups, and visualized as pie charts in **Figure S5**. Similar to the level 67 results, the majority of the phospho-ERK1/2^+^ perikaryon profiles in the DMX at level 69 were identified to be ChAT^+^ and negatve for DBH in both saline- (99.7%) and 2-DG-treated (100%) subjects (**Fig. S5**, pERK+ row, pERK1/2^+^ & ChAT^+^; *magenta-colored* slice). However, among total ChAT^+^ perikaryon profiles, the percentage of pERK1/2^+^ & ChAT^+^ double-positive perikaryon profiles appears to be greater in 2-DG- (53.5%) than saline-treated (43.6%) subjects, but the difference is not as prominent as for the level 67 relative percentages.

## 4. Discussion

In this study, we found that an intravenous glycemic challenge, 2-deoxy-D-glucose (2-DG), is associated with the rapid activation of *rhombic brain (His, 1893)* (RB) [21] neurons, specifically those in the *locus ceruleus (Wenzel & Wenzel, 1812)* (LC) [83] and *nucleus of solitary tract (>1840)* (NTS). These were sampled and mapped to discrete rostrocaudal levels of the *Brain Maps 4.0* open-access rat brain atlas [82]. The activation was encoded by elevations in the numbers of perikaryon profiles displaying immunoreactivity for the phosphorylated forms of ERK1/2 in these regions within 15 minutes of the challenge. These activated neurons included catecholaminergic and non-catecholaminergic neurons, some of which also displayed basal levels of activation in saline-treated subjects. Below, we discuss these findings in relation to the relevant physiology and anatomy of neural circuits engaged during feeding and glycemic counterregulation.

### 4.1 Physiological considerations

The activation of neural circuits during glycemic challenge, as with other systems-level processes, can be observed and measured at multiple scales, with careful considerations across these scales providing a convergence of evidence in support of the conclusions one draws for the whole system. Here, we consider multiscale questions for our measured readout (phospho-ERK1/2 immunoreactivity), our chosen stimulus (2-DG injection), and their coupling in the brain, with an eye towards an accounting of how these components, when considered together, provide evidence that the brain is mounting a physiological response to a glycemic challenge, a response that has spatiotemporal components that can be reliably tracked.

#### 4.1.1 Phospho-ERK1/2 as rapid trackers of neuronal activation

As with most phenomenological concepts in science (pp 3–5 of [114]), the term ‘activation’ means different things under different contexts. In the brain, activation could refer to any of myriads of changes in structural or functional conditions, including, to name just a few: neuronal firing rates, intracellular Ca^2+^ metabolism, neurotransmission, gene transcription, protein translation or post-translational modifications. Whether such measures of activation always reflect *relevant* physiological changes underlying a given function or set of functions has been the subject of much discussion [78, 79, 115–120].

Here, we refer to ‘activation’ as a change in the phosphorylation status of mitogen-activated protein (MAP) kinases in RB neurons in association with 2-DG treatment. MAP kinases ([EC 2.7.11.24](https://iubmb.qmul.ac.uk/enzyme/EC2/7/11/24.html)) comprise protein kinases catalyzing a hierarchical, multi-tiered regulatory cascade of phosphorylation events [121]. Components of the ERK subfamily of MAP kinases have been localized within the RB structures we examined here, including the LC, NTS and DMX [122]. Phosphorylation of ERK1/2 kinases at residues within a Thr–X^aa^–Tyr dual-phosphorylation motif in the enzyme activation loop (in human: Thr^202^ and Tyr^204^; in rat: Thr^183^ and Tyr^185^) has been demonstrated to be essential for catalytic activity [123]. The use of phospho-specific antibodies against MAP kinase is an established method of indirectly identifying a catalytically active enzyme [124], although the phosphorylated residues bound and detected by these antibodies are, technically, products of phosphorylation reactions catalyzed by the MAP kinases hierarchically upstream of the immunolabeled enzyme [124].

Importantly, although we define activation here in terms of phosphorylation state changes, we do not measure phosphorylation *per se*, but the numbers of neurons displaying changes in immunoreactive signal for phosphorylated MAP kinases. The phospho-specific antibodies are selective for labeling only the phosphorylated and not the unphosphorylated forms of the enzyme. Thus, this labeling reflects activation of a pool of MAP kinase molecules, as observed from accumulated immunoreactive signals in the perikaryon, and at times the nucleus, of the imaged cell profile. The resulting immunoreactive signal, which varied in intensity between saline- and 2-DG-treated subjects and across regions, was not quantitatively compared here. Rather, the numbers of perikaryon profiles displaying immunoreactive signal above background levels were counted as a means to assess population-level recruitment of cells.

We observed greater numbers of phospho-ERK1/2^+^ neuronal profiles in the LC at level 51 and NTSm at levels 67 and 69 in 2-DG-treated subjects relative to saline-treated control subjects. The physiological significance of this alteration in the numbers of neurons immunoreactive for phospho-ERK1/2 is not known at this time. However, we infer from these changes that greater numbers of neurons are being ‘recruited’ to perform physiologically-relevant activities in the face of this glycemic challenge. In support of this view, three lines of evidence suggest that the increases reflect a change in the overall signaling status of LC and NTSm neurons to initiate *de novo* changes in gene expression and/or changes in neuronal firing rates in response to 2-DG administration. These sets of evidence will be considered in order of their decreasing relevance to the context of glycemic challenge.

First, we have previously shown that, in association with glycemic challenges such as intravenous delivery of insulin or 2-DG, phospho-ERK1/2 reliably tracks cellular activation within the *paraventricular hypothalamic nucleus (>1840)* (PVH) and *arcuate hypothalamic nucleus (>1840)* [66–69, 71]. Furthermore, under these conditions, at least where we have established it for the *parvicellular division (>1840)* of the PVH, phospho-ERK1/2 not only tracks cellular activation during these challenges, but likely plays a causal role in downstream processes engaged by these challenges, such as recruitment of transcription factors, induction of certain gene expression patterns, and increases in neuronal firing rates [69, 72]. Thus, it seems reasonable to consider the possibility that, under the same glycemic challenges, the phospho-ERK1/2 observed in RB neurons may also serve similar functions. However, the links in the chain of evidence required to establish a causal role for phospho-ERK1/2 in RB neuronal responses to glycemic challenge, as it has been established for PVH responses to these challenges [70], have yet to be forged.

Second, outside of our studies of phospho-ERK1/2 in association with glycemic challenge in the RB, these enzymes have been studied under other contexts in this subdivision of the brain. For example, in the LC, ERK1/2 tracks cellular activation associated with pain-related anxiety in male arthritic rats [125] and appears to contribute to tonic inhibition of LC neurons under certain experimental conditions [126]. Optogenetic stimulation of hypocretin/orexin-expressing neurons in the hypothalamus triggers elevations in biochemically detectable phospho-ERK1 and 2 in LC tissue homogenates [127]. In the NTS, phospho-ERK1/2 mediates the suppressive effects of cholecystokinin or a melanocortin receptor agonist on food intake [128, 129]. In other RB regions, including those that correspond to the functionally-defined rostroventrolateral medulla, the ERK pathway is involved in urotensin-induced increases in sympathetic vasomotor tone [130]. Thus, throughout portions of the RB, phosphorylated ERK1/2 appear to participate within causal chains of signal transduction that link stimuli to appropriate physiological responses. This line of evidence, therefore, supports a similar role for ERK1/2 under conditions of glycemic challenge.

Finally, the more general question of whether phospho-ERK1/2 elevations correlate with other increases in ‘activation’, such as elevated firing rates for neurons or increased signaling cascades leading to gene expression, has been systematically addressed by Fields and colleagues in a number of incisive studies using dorsal root ganglion neurons as a model system. From these studies, they have established that ERK1/2 is tuned to track specific patterns of action potential firing with a fairly high fidelity and coordinates transcriptional networks [131–134].

Taken together, these three lines of evidence offer us with independent but conceptually integrable sets of experimental results suggesting that the phosphoryl modifications of ERK1/2 kinases, under these experimental conditions, constitute an intracellular biochemical process that rapidly and reliably tracks our intervention. Moreover, they support the notion that not only are they tracking our intervention, they could be involved in mediating the physiologically-meaningful responses to it. Having established this framework for our measured response, it is the nature of the intervention we used which concerns us next.

#### 4.1.2 Intravenous 2-deoxy-D-glucose as a physiologically-relevant glycemic challenge

While phospho-ERK1/2 is well-positioned as a reliable readout of glycemic challenge, is 2-DG a physiologically-relevant stimulus for producing such a challenge? Historically, Warburg’s discovery of aerobic glycolysis in tumors [135, 136] contributed to the great interest in utilizing glycolytic inhibitors, such as 2-deoxy-D-glucose (2-DG), as a means to impair tumor glucose utilization [*e.g*., 137, 138], an interest that led to its use as a treatment for cancer patients [139]. This strategy of impairing glucose utilization also found currency in other clinical domains, most notably in using 2-DG to assess physiological counterregulatory responses in humans [140] and other animals [141–143]. At the scale of the intact living organism, 2-DG has also been used as a means to experimentally induce food intake [73, 142–148], a behavioral response that is influenced by time of day [146], diet [147], and nutritional status [148]. Notwithstanding that feeding and glucoregulatory control circuits are likely components of overlapping but nonredundant neural systems, the effects of 2-DG on both counterregulatory processes and food intake have been established well enough to provide justification of its continued utility for exploring these systems at this scale.

This assertion is further supported by studies of 2-DG that have been performed at the cellular or cell-free (isolated biochemical) scales. In particular, enzyme kinetics studies, each conducted under distinct experimental starting conditions, provide a striking convergence of circumstantial or direct evidence that 2-DG is an inhibitor of glycolysis [149–152]. The main nodal points for direct 2-DG binding –– and the inhibition of glycolysis it causes –– appear to be at reaction steps catalyzed by hexokinase ([EC 2.7.1.1](https://iubmb.qmul.ac.uk/enzyme/EC2/7/1/1.html)) [149, 151, 152] and glucose-6-phosphate isomerase (GPI; [EC 5.3.1.9](https://iubmb.qmul.ac.uk/enzyme/EC5/0301p.html#0109)) [150]. Although 2-DG has been empirically determined to bind with a lower affinity to brain hexokinase than glucose does (in rat: K_m_ = 1.1 × 10^−4^ mol/L for 2-DG vs. 4.5 × 10^−5^ mol/L for glucose [153]), Lineweaver-Burk [154] analyses applied to assay measurements of hexokinase kinetics revealed that 2-DG serves as a competitive inhibitor of the enzyme in the presence of varying concentrations of glucose, but not Mg-ATP [151]. Kidney-derived GPI, in contrast, has been shown in a cell-free system to be a principal target of 2-DG-6-PO_4_ [150], which is produced by hexokinase in the presence of 2-DG. Taken together, the data indicate that there are multiple primary modes to block glycolysis by 2-DG. Moreover, allosteric feedback inhibition of hexokinase by 2-DG-6-PO_4_ could serve as a secondary block of the pathway [150]. Targeted inhibition at these steps appears to be reflected in the disposition of glucose utilization *in vivo* [155], and hexokinase isoforms of the glucokinase family ([EC 2.7.1.2](https://iubmb.qmul.ac.uk/enzyme/EC2/0701a.html#2)) are expressed in the brain, and have recently been found to be critical for glucoprivic feeding triggered by 2-DG [156]. Thus, biochemical and physiological evaluations of 2-DG actions support its role in impairing glucose utilization.

Yet, despite the evidence for its direct role as a glycolysis inhibitor, evidence has also been gradually accumulating for more complex actions of 2-DG that occur alongside and perhaps as a consequence of its actions on intermediary metabolism. Early work using isolated peripheral nerve preparations demonstrated an inhibitory effect of 2-DG on electrogenic activity of the sodium-potassium pump in firing nerve [157, 158], and 2-DG appears to have observable effects on neural electrical activity more centrally, by triggering extrasynaptic GABA release which results in tonic GABA currents in central neurons at the network level, and inhibition of K_ATP_ channels in single neurons [159]. These observations have important clinical implications for individuals who are placed on a ketogenic diet to treat seizures [160]. Thus, aside from and/or perhaps as a result of 2-DG actions on glycolysis, neuronal and circuit-level electrophysiology is altered in the brain. Considered together with recently reported observations that brain endothelial cells undergo aerobic glycolysis (as described above for tumors) and that 2-DG can alter blood-brain-barrier transcellular transport mechanisms and permeability in these cells [161], it would appear that the effects of systemic 2-DG injection are diverse and complex. While here we have briefly just enumerated those effects that still fall within the contexts of glycemic challenge and inhibition of metabolism, we now consider below effects that appear somewhat distinct from these contexts.

#### 4.1.3 Stimulus-response coupling: 2-DG coupling with phospho-ERK1/2 and alternate coupling relationships and associations

In this study, we provide evidence of an association between 2-DG administration and increases in the numbers of observable cells in our sample that display immunoreactive phospho-ERK1/2. There are compelling reasons, discussed above, supporting the physiological relevance of using 2-DG as a glycemic challenge and the reliability of the ensuing phospho-ERK1/2 responses to track this challenge. Nevertheless, invoking cause-effect relationships between external stimuli and intracellular signaling cascades is challenging (*e.g*., see Ref. [162]), and the linkage between the two in our study awaits further strengthening and should be considered here as an initial characterization of a stimulus-response relationship. Many questions remain. 2-DG, for example, is a mannose, and as such, it can profoundly affect cells by altering their complement of glycosylated proteins [163, 164], which could alter the sensitivity or responses of cells being studied here. Thus, 2-DG could be coupled to glycosylation signaling processes independent of its glycolytic targets or downstream effects on electrical signaling or blood-brain-barrier permeability. 2-DG can also trigger a mild hypothermia and hypotension when given systemically [165, 166], responses we did not measure in our study. It is not clear if these physiological changes engage the same *rhombic brain (His, 1893)* [21] (RB) neuronal populations that glycemic challenge does, although one laboratory has insightfully examined the populations engaged by hypotension and glucoprivation ([166]; see *Section 4.2* for more on this).

Along similar lines, the argument that 2-DG can trigger multiple physiological conditions finds a counterpoint from our observations, thus far reported only in preliminary form [85, 87], that phospho-ERK1/2 can be triggered in the same RB regions by multiple glycemic challenges. Specifically, we have reported that intravenous injection of insulin can trigger phospho-ERK1/2 in the LC [85] much as 2-DG does in the present study. These findings provide compelling support for the idea that the coupling of 2-DG to phospho-ERK1/2 is encoding a physiological response to changes in glycemic status. Our work also builds upon earlier work conducted by the Watts laboratory demonstrating that numbers of phospho-ERK1/2-immunoreactive neurons are elevated in the NTS and DMX in association with low concentrations of blood glucose and also exhibit a time-of-day difference in peak expression [68]. Thus, phospho-ERK1/2 does appear to track closely with glycemic challenges in portions of the RB, although it is now clear that there are other key molecules that may contribute to glucosensing or mark the presence of glucosensing neurons (*e.g*., AMP kinase [167, 168]; GLUT2 transporter [28]; phospho-TH [169]). The activation of these molecules is likely only salient in specific places in the RB and under varying conditions, encoding glycemic and other processes in a manner germane to the subsystems the molecules serve. One could imagine that they essentially represent context-specific combinatorial codes – an emergent “biodigital jazz” [170] that allows a dynamic encoding of ever-changing events (also see Refs. [79, 133, 171]). We now discuss the physiological importance of such encoding in the RB in relation to our results.

#### 4.1.4 Physiological significance

Keeping in mind the previous discussion concerning the reliability of our measured readout, the relevance of 2-DG as a glycemic challenge, and how well the two might be coupled, we turn now to consider the functional importance of our results. First, we consider the rapid recruitment of LC neurons observed after 2-DG challenge. Given the role the LC plays in controlling wakefulness, arousal and cognition (*e.g*., Refs. [172, 173]), this recruitment can be integrated conceptually within the broader framework of producing or resetting arousal state during glycemic challenge, perhaps as a means to assist the organism in awareness of nutrient and/or energy deficit. If true, this may reflect a recruitment of a set of LC-connected circuits that allows the organism to mount a larger-scale resetting of behavioral state, in parallel to the circuits transmitting signals encoding glucosensing and counterregulatory responses being handled also or exclusively by other RB structures, such as the NTS and DMX. Such functional dichotomy may help explain observations that hypoglycemia unawareness can be reversed in insulin-dependent diabetics without effectively reversing impaired glucose counterregulation [174]. Perhaps the neural systems producing these distinct clinical signs of energy deficit are also distinct, or at least partly so. It is clear, however, from recent evidence for a modular organization of neural elements in the LC [175, 176] that this structure is well-suited to handling both sets of information to effect both outcomes. That the LC can receive neurally-encoded information about glucose deficit is supported by electrophysiological evidence, in the mouse, that neurons in the rostral ventrolateral medulla, known to be activated by hypoglycemia and many other physiological conditions, can monosynaptically activate LC neurons by releasing glutamate [177]. Moreover, cellular activation in the LC has been tracked using Fos expression following systemic or fourth ventricular injection of insulin or 2-DG [36, 39, 178–180], and glycemic challenges are also associated with altered LC single-unit activity [181], tyrosine hydroxylase expression [182] and phosphorylation [183] (but see Ref. [169]), and norepinephrine transporter expression [184]. Insulin administration is also associated with decreased glucose utilization in the LC, but evidence indicates that this effect may be a direct effect of insulin itself and not the hypoglycemia produced by insulin [185, 186].

However, the LC also figures prominently in a bevy of studies demonstrating that its neurons are highly sensitive to perturbations caused by hypoxia or hypercapnia, adjusting their excitability to conserve energy under such conditions (*e.g*., [187–190]). The literature therefore indicates that LC neurons may be multimodally responsive to a variety of physiological perturbations, not just those that affect glycemic status (see also Ref. [180]). The elevated numbers of neurons immunolabeled for phospho-ERK1/2 in the LC of 2-DG-treated subjects may reflect a generalized mechanism of activation for the neurons of this structure, one that is not necessarily encoding signals *exclusive* to changes in glycemic status.

This may also be true for the NTSm. Like cells in the LC, cells within this structure appear to be responsive to hypoxia and hypercapnia [186]. On the other hand, much evidence also indicates that NTS neurons actively respond to changes in glycemic status [23–29, 31, 192, 193]. They possess ATP-sensitive potassium channels which appear to confer glucosensing capabilities to them [192]. These neurons depolarize in response to low extracellular glucose, but not in the presence of high glucose [193]. Thus, the phospho-ERK1/2-immunoreactivity we observed in NTS perikaryon profiles may reflect activation of glucosensing elements. Some of these profiles did not display immunoreactivities for either ChAT or DBH in the NTS and are therefore members of one or more additional phenotypes that are known to populate this region [194]. It is also unclear if these are neuronal profiles; indeed, the NTS has astrocytes critical for glucosensing, counterregulatory responses and other functions [26, 27, 49] (reviewed in Ref. [195, 196]).

Finally, as we close out our discussion of functional considerations concerning our findings, sev-eral open questions remain about the roles of these activated regions for larger processes of metabolic regulation and feeding control. For example, since the exact organization of the extended networks served by these activated regions remains unknown at the present time, it remains unclear whether there are subsystems that allow for the body to adequately switch its anaerobic versus oxidative capacity [136]. Nor is it clear whether this switch in metabolism in the face of glycemic challenge allows the organism to switch macronutrient diets accordingly [45] and to tailor its ability to counterregulate and/or ingest additional food to gain energy.

### 4.2 Anatomical considerations

In this study, we identified the LC at level 51, and the NTSm and DMX at levels 67 and 69, as harboring elevated numbers of phospho-ERK1/2-immunoreactive neurons, which were identified as being either catecholaminergic or non-catecholaminergic; these neurons have been implicated in maintaining arousal state, autonomic control and/or nutrient sensing [197, 198]. Below, we discuss these principal findings in relation to our methodologies and also in relation to the known anatomy of the RB.

#### 4.2.1 Standardized mapping of activation patterns in the brain

A key goal of this study was to build upon and expand the prevailing literature by explicitly mapping patterns of neuronal activation to glycemic challenge using a standardized rat brain atlas. As we have argued previously [80, 98, 199, 200], the use of a standardized atlas to formally map cellular distributions affords investigators with the ability to examine multiple spatial datasets in a form that facilitates their direct comparison. We opted, as with all of our spatial mapping efforts for the rat (*e.g*., Refs. [98, 199, 201]), to utilize the open-access rat brain atlas of Swanson [82], which is accompanied by a defined nomenclature that is traceable to the primary literature.

By mapping the patterns of immunoreactivity that we observed to this reference space, we can align and interrelate our datasets with those of our own and of other laboratories that have mapped their datasets to the same atlas. For example, the Watts laboratory has shown [68] that at plasma glucose levels < 3.15 mM, increased numbers of phospho-ERK1/2-immunoreactive cells are present within the NTS and the DMX at rostrocaudal locations that fall within levels 65–66, 68–69, and 69–70 of the *Brain Maps 4.0* atlas. Importantly, at plasma glucose levels > 3.15 mM, the numbers of cells that display such immunoreactivity were much fewer by comparison. In the present study, 2-DG was utilized as the glycemic challenge, which increases plasma glucose levels while decreasing glucose utilization. Consistent with the findings of the Watts laboratory, then, the activation we observed at these atlas levels may be attributable to decreases in glucose utilization rather than hyperglycemia *per se*.

Moreover, other activation has been noted and mapped at or near to the atlas levels we report here. For example, peripheral injection of the GLP-1 receptor agonist, exendin-4, triggers Fos expression in the rat LC and NTSm, in neuronal populations mapped to atlas levels 52, 66, 70, and 71 [202]. On those same levels, GLP-1 immunoreactive cells and fibers were also mapped and localized to NTS subregions and surrounding areas that were largely non-overlapping with the Fos-immunoreactive neurons. Similarly, we observed increases in Fos-immunoreactive NTSm neurons (and notably, virtually no DMX neurons) at atlas level 69 in animals allowed to feed for 2 hours after an imposed 40 hr fast, as compared to subjects that were fasted for 40 hrs but not allowed to feed again [98]. This level of the NTSm is the same level in the present study where we observed neurons to be recruited by our glycemic challenge. Thus, data gathered by our laboratory from two separate studies indicate that the NTSm, at atlas level 69, contains neurons responsive to refeeding or glycemic challenge. It remains to be determined whether the same neurons can be activated in this region by both perturbations.

##### 4.2.2 Alignment of activation patterns with connectivity data in the brain

The patterns of activation that we document here can also be placed in register with known neural connection patterns established by neuroanatomical tract-tracing experiments that have been mapped using the *Brain Maps 4.0* [82] framework or its earlier iterations. For example, the rat LC at level 51 reportedly receives input from a brain region, designated as MnPO by the authors [203], which corresponds essentially identically with the *median preoptic nucleus (Loo, 1931)* [204] (MEPO) in *Brain Maps 4.0* [82]. Using the *BM4.0* framework, these investigators carefully charted the axonal boutons and terminal fields of anterogradely-labeled fibers from injections of 10-kDa biotinylated dextran amine (BDA) [205] into this structure. Clearly-identifiable axons and boutons at the light-microscope level were observed in the LC (see Fig. 7a–c in Ref. [203]), which were found to closely appose tyrosine hydroxylase-immunoreactive cells of the LC (see their Fig. 7a,b). These projections were displayed on digital templates accompanying the second edition of *Brain Maps* [206], and their existence was confirmed by injection of a retrograde tracer into the LC. The retrograde pattern of labeling, interestingly, revealed labeled cells not only in the MEPO, but also in extrahypothalamic regions, including the *bed nuclei of terminal stria, anterior division, oval nucleus (Ju & Swanson, 1989)* [207] (BSTov; see their Fig. 8f’), a region which has been previously reported in preliminary communications [79, 208, 209] as also displaying elevations of phospho-ERK1/2-immunoreactive neurons following 2-DG administration. Collectively, these nascent observations would indicate that the atlas-based mapping of MEPO→LC and BSTov→LC connections, together with our atlas-based mapping of BSTov [79, 208, 209] and LC activation (Refs. [85–90]; the present study) in association with 2-DG treatment, reveals a larger set of neural connections that may participate in relaying descending information to the LC that could be salient during glycemic challenge. This is also supported by evidence for a set of connections to the LC, albeit sparse, that have been reported from the *lateral hypothalamic area, juxtaventromedial region, dorsal zone (Swanson, 2004)* [210] LHAjvd [211]. These connections, however, invest the NTS more heavily, as shown in maps of regions of the NTS slightly more caudal to the level we mapped here (see their Fig. 4, panels AAA and BBB).

Although the connections between peripheral organs and neuronal populations in the RB have been intensively studied, few have traced and mapped these connections in the rat using the *Brain Maps* atlas framework in its various editions. However, Garcia-Luna *et al*. [76] have provided an elegant demonstration of the power of atlas-based mapping using this spatial model to help register peripheral nervous system connections with neuronal populations in the RB. Specifically, these colleagues have shown that vagal sensory endings innervating the wall of the hepatic portal and superior mesenteric veins make multi-order connections with neuronal populations in the *dorsal motor nucleus of vagus nerve (>1840)* (DMX). Furthermore, they showed that injections of viral tracer into the left nodose ganglion of rats produced transsynaptic labeling that, as a function of survival time, progressively reached NTS glial cells (24 h), and AP, NTS and DMX neurons (48 and 72 h).

#### 4.3 Limitations of this study

In this study, we provided new mapping and atlas-based localization of neuronal populations activated within 15 minutes of an intravenous delivery of a glycemic challenge. This activation is defined in this study as the elevations in the numbers of perikarya (cell body) profiles immunoreactive for phospho-ERK1/2. A few caveats must be kept in mind regarding the use of this marker to track activation. First, unlike in other systems [69, 70, 72, 128, 130, 212], phospho-ERK1/2 has not yet been established to be within the causal chain of events that link systemic glycemic challenge to hypoglycemic counterregulation at the level of RB. Second, it is not clear whether the activation observed is a result of neural inputs triggering network activity, and/or results from a direct action of 2-DG on the cells themselves. The latter possibility is certainly plausible, considering that peak accumulation of [^14^C]2-DG in rat brain tissues is observed in as little as five minutes following its intravenous administration (*e.g*., see Fig. 5 in Ref. [213]). Finally, only a single time point was examined in this study, and this, only in very circumscribed portions of the LC, NTS and DMX. High-throughput cell counting methods involving deep learning algorithms and computer vision, such as those that we are developing [214], could streamline analytical workflow procedures to allow for a broader spatiotemporal survey of brain-wide activation patterns following glycemic challenge.

## 5. Conclusions

The results of this study provide new evidence that that there are rapidly activating neurons, and perhaps glial elements, that are engaged within 15 minutes of glycemic challenge in neurons of the LC and NTSm. Some but not all of these neurons are catecholaminergic. Importantly, we mapped these neurons to standardized atlas templates from *Brain Maps 4.0*, an open-access rat brain atlas that allows for data from multiple labs to be compared with one another. We discuss how our datasets match up with those showing neural connections atlas-mapped to these regions from the hypothalamus or periphery. The new maps furnished in this study constitute an important step towards a global brain-wide assessment of cellular responses to glycemic challenges, which may allow for more refined clinical or therapeutic interventions for those individuals experiencing complications as a result of iatrogenic hypoglycemia.

## Supporting information

Supplemental Methods, Figures, Tables and References

## Author Contributions

Experimental conception and design: AMK. Project management: AMK, RHT, GPT, SB. Surgeries and *in vivo* experiments: LJA, AMK. Perfusions and histology: LJA, AA/SDC, GS. Tissue inventory: AMK, GPT, AA, SDC. Immunohistochemistry: SDC, VIN. Microscopy and imaging: GPT, AMK, SDC. Cytoarchitectonic analysis and atlas-based mapping: GPT, JSP. Quantitative and exploratory data analyses: SB, GPT, JSP, AMK. Artboard construction: AMK, GPT, VIN, SB. Research funding procurement: AMK, RHT. Research supervision and training: AMK, GPT, RHT, SB, LJA, VIN. Conference presenters: GPT, JSP, SDC, AA/ GS. Manuscript preparation: AMK, GPT, SB, VIN with final checks from all co-authors. Manuscript revisions: AMK, GPT. All authors have read and agreed to the published version of the manuscript.

## Institutional Review Board Statement

The animal study protocol was approved by the Institutional Animal Care and Use Committee of The University of Southern California (protocol code 11249 approved on August 26, 2009).

## Data Availability Statement

All relevant protocols and data are contained within the article, along with its Supplemental Files.

## Acknowledgments

We are grateful to Dr. Alan G. Watts and Graciela Sanchez-Watts (USC) for allowing the generous use of their surgical and laboratory spaces during the early stages of this project, Dr. Larry W. Swanson (USC) and Cathleen Crayton for facilitating weekly discussions of our group in Dr. Swanson’s research space and for the use of his laboratory for tissue processing, and Dr. Ted Berger (USC) for use of his laboratory. Tiffanie Nham, Michael Zobel, and Jordan Michaels participated in valuable discussions. Jeannie Y. Zhang, Danielle Goodrich, Michael Zobel, Ellen M. Walker, Evangelina Espinoza, Kenichiro Negishi and Marina Peveto provided valuable technical assistance or datasets, and Dr. Anne Jokiaho (USC) graciously facilitated access to legacy records at USC. AMK and RHT thank Dr. David Glasgow in the USC Undergraduate Education Office for his support of our students. We also thank Dr. Armando Varela-Ramirez for his assistance in the Cellular Characterization and Biorepository Core Facility of the UTEP Border Biomedical Research Center and Dr. Larry W. Swanson for clarifications concerning his atlas nomenclature. All *BM4.0* atlas maps were modified and/or reproduced with permission under the conditions outlined by a Creative Commons BY-NC 4.0 license. The authors express their appreciation for the valuable digital resources provided by the Internet Archive and The University and State Library of Saxony-Anhalt in Halle, Germany. AMK and GPT dedicate this study to the memory of the late Dr. Sue Ritter (Washington State University), for her pioneering and systematic investigations of hindbrain mechanisms of glucoregulatory control. AMK thanks his friends at DSB for their support during the final stages of this project.

## Abbreviations

AP: *Area postrema (>1840)*
DMX: *Dorsal motor nucleus of vagus nerve (>1840)*
LC: *Locus ceruleus (Wenzel & Wenzel, 1812)*
LDT: *Laterodorsal tegmental nucleus (>1840)*
MEV: *Midbrain nucleus of trigeminal (>1840)*
MY: *Medulla (Winslow, 1733)*
NTS: *Nucleus of solitary tract (>1840)*
NTSce: *Nucleus of solitary tract, central subzone (>1840)*
NTSco: *Nucleus of solitary tract, commissural part (>1840)*
NTSge: *Nucleus of solitary tract, gelatinous subzone (>1840)*
NTSl: *Nucleus of solitary tract, Lateral part (>1840)*
xsNTSm: *Nucleus of solitary tract, Medial part (>1840)*
NTSmc: *Nucleus of solitary tract, Medial part, Caudal zone general, caudal subzone (>1840)*
NTSmcg: *Nucleus of solitary tract, Medial part, Caudal zone general (>1840)*
PB: *Parabrachial nucleus (>1840)*
PCG: *Pontine central gray (>1840)*
PCGg: *Pontine central gray general (Swanson, 2004)*
PGRN: *Paragigantocellular reticular nucleus (>1840)*
PGRNl: *Paragigantocellular reticular nucleus, Lateral part (>1840)*
RB: *Rhombic brain (His, 1893)*
scp: *superior cerebellar peduncle (Procháska, 1800)*
ts: *solitary tract (>1840)*
V4f: *Floor of fourth ventricle (Reil, 1807–1808)*
XII: *Hypoglossal nucleus (>1840)*
z: *Nucleus z (>1840)*

