## Supplemental Methods, Figures, Tables and References for "Glycemic challenge is associated with the rapid cellular activation of the locus ceruleus and nucleus of solitary tract: Circumscribed spatial analysis of phosphorylated MAP kinase immunoreactivity"

**Supplemental Materials: Tapia et al.**

### A. Supplemental Methods with References

***1. Intra-atrial catheterizations.*** The throat was shaved, disinfected with surgical scrub, and the right jugular vein located under the guidance of a dissecting stereoscope and fiber-optic illumination. The surgical target area harboring the vein was apparent under the microscope by the fluttering of the overlying skin with each heartbeat. Once the area was located in this manner, cold-sterilized forceps and surgical scissors were used to lift up the skin and make an incision to expose the jugular vein. A previously-prepared cold-sterilized catheter assembly consisting of silastic tubing (0.020" i.d., 0.037" o.d.; Catalog # 76P020-210-037; Specialty Manufacturing, Inc., Saginaw, MI), held fast by cured cyanoacrylate glue and a 23-gauge stainless steel connector to larger-diameter Tygon tubing, was first flushed with sterile 0.9% saline, and then inserted slowly through the vein and into the atrium of the heart. It was held firmly but loosely in place with two lines of suture filament threaded under the vein and catheter assembly. Once secured in this manner, the catheter was gently aspirated using a 1.0 ml syringe to ensure blood flow from the heart, and the filaments untied if adjustments were needed. Once patency was confirmed, the filaments were then tied securely, as were two additional filaments, and the larger end of the tubing was exteriorized by tunneling it under the skin and out through the subscapular space. The entire procedure took approximately 15–20 min for each animal, after which the incision was closed using wound clips and cleaned.

***2. Antibody information.*** Preadsorption experiments performed by ourselves or other laboratories have demonstrated the specificity of the primary antibodies we used in this study for binding to their intended antigen targets. First, as noted previously [1], preadsorption of the rabbit phospho-ERK1/2 antibody we used in this study with the immunizing peptide used to produce it abolished all immunostaining in brain tissue (AMK; *unpublished data*). Second, the mouse monoclonal anti-DBH antibody we used reportedly produces immunoreactivity that is completely abolished in rat brain tissue sections when the antibody is preadsorbed with a 10-fold excess of bovine adrenal DBH [2]. Similarly, the primary anti-ChAT antibody we used, which was raised against human placental ChAT, reportedly did not produce immunoreactive signal when preadsorbed with a rat recombinant ChAT protein [3]. Additionally, Sutton and colleagues [4, 5] have demonstrated that rat *nucleus of solitary tract (>1840)* (NTS) tissue displays specific bands for phospho-ERK1/2 using the same antibody we used in our study, bands which can be co-labeled using an antibody against the unphosphorylated forms of the enzyme.

#### *3. Specificity controls for immunohistochemistry*

*3.1 Control experiments to test for secondary antibody cross-reactivity.* A series of tests were performed to check the specificity of the secondary antibodies we used for their primary antibody targets. First, using a single series from saline- (#K10-011) and 2-DG-treated (#K10-026) subjects, we performed immunofluorescence histochemical experiments as noted above, but using secondary antibodies that were purposefully mismatched to the original primary antibodies. Thus, immunofluorescent staining for DBH was conducted using sections containing the *nucleus of solitary tract (>1840)* (NTS) (to ensure DBH was present) selected from the saline-treated animal case. Visual confirmation of the presence of the NTS was performed using surrounding white matter landmarks made visible under dark-field illumination (Olympus BX63 microscope, ×10 magnification objective lens; *see Section 2.6.1 for details*). The primary mouse anti-DBH antibody was used in conjunction with both a Cy3-labeled donkey anti-rabbit IgG, and an Alexa 647-conjugated donkey anti-goat IgG. The Alexa 488-conjugated streptavidin reagent was also added to the reaction. The primary antibody incubations lasted 18 h at 4 °C, the secondary antibody incubations for 5 h at room temperature, and the incubation of Alexa 488-conjugated streptavidin for 1 h at room temperature. Similar experiments were conducted for anti-pERK1/2, anti-DBH, and anti-ChAT using selected sections containing the *nucleus of the solitary tract (>1840)* (NTS), *locus ceruleus (Wenzel & Wenzel, 1812)* [6] (LC), *dorsal motor nucleus of vagus nerve (>1840)* (DMX), and *hypoglossal nucleus (>1840)* (XII) from a 2-DG-treated subject (#K10-026).

*3.2 Control experiments to test for nonspecific binding of secondary antibodies to tissue sections.* To ensure that secondary antibodies were not producing labeling patterns as a result of a direct and non-specific binding to endogenous proteins in the tissue, we performed our fluorescence immunohistochemistry on a subset of tissue sections for which the primary antibodies were not included in the reaction sequence. To keep consistency across controls, we selected tissue sections from a 2-DG treated subject (#K10-026) that was comparable to those selected for the saline-treated subject tested in *Section 2.4.2.3*. The secondary antibodies were reacted as a cocktail, and consisted of the Cy3-labeled donkey anti-rabbit IgG, the biotinylated donkey anti-mouse IgG, and the Alexa 647-labeled donkey anti-goat IgG listed in **Table 1**. Sections were incubated at 4°C for 18 h in blocking solution (to mimic the reaction duration ordinarily used for the primary antibody step) and 5 h in the secondary antibody cocktail at room temperature.

*3.3 Checks for tissue autofluorescence.* To verify that unlabeled tissue sections did not exhibit any endogenous fluorescence contributing to the signal observed, sections containing the *locus ceruleus (Wenzel & Wenzel, 1812)* [83] (LC) were first selected by carefully checking for tissue sections containing the structure as observed under dark-field illumination. Cryoprotectant was removed from sections in a series of 5 × 5 washes in TBS in a netted well tray. Sections were removed from the netted well tray, mounted onto Superfrost™ slides, and allowed to air-dry for 1 h under light-protected conditions. The slides were coverslipped using bicarbonate-buffered glycerol mounting solution and sealed with clear nail polish around the edges of the slide. Photomicrographs were then taken using an Olympus BX63 microscope at ×10 and ×20 magnifications to check for observable signal in the tissue.

*3.4 Checks for binding of Streptavidin-Alexa 488 to endogenous biotin in tissue.* We aimed to account for any endogenous binding of streptavidin used to stain the biotinylated secondary antibodies used in this study. As before, sections containing the LC were selected under dark-field illumination, rinsed of cryoprotectant with TBS (5 × 5 washes), and transferred to a 24-well plate containing Alexa 488-conjugated streptavidin solution for 1 h at room temperature. Sections were then rinsed again (5 × 5) in TBS, mounted onto Superfrost™ slides, and allowed to dry for 1 h. Slides were as before coverslipped and sealed with nail polish. Photomicrographs were again taken using an Olympus BX63 microscope at ×10 and ×20 magnifications to check for observable signal in the tissue.

### *4. Additional details concerning confocal microscopy*

*4.1 Confocal scanning.* Under a *Continuous* scan mode, we used the lookup table *Range Indicator* in the software to guide *Gain (Master)* and *Digital Offset* adjustments to keep the image signal spread over the dynamic range of the detector, and to prevent missing pixels or pixel saturation as a result of under- or overexposed regions of tissue, respectively.

*4.2 Visualization of far-red fluorescent signals.* For preparation of certain photomicrographs of ChAT-immunolabeled cells in the far-red channel, which were acquired from tissue displaying a faded fluorescence, brightness and contrast were enhanced in Adobe Photoshop to help the reader visually differentiate the faint cells from background and to aid the reader in assessing patterns in two-channel images; these changes, which were applied to the whole image in each instance, are noted in the caption of each relevant figure. For these images, along with the remaining images shown in this study, care was taken that all acquisition and post-acquisition parameters were identical for images displayed together from control and 2-DG treatment groups, in keeping with community standards (*e.g.*, [7]).

### *5. Checks for fluorescence cross-talk*

*5.1 Evaluating crosstalk at the level of fluorescence labeling.* Three approaches were employed – two precautionary and one empirical – to help exclude the possibility that secondary antibody-fluorophore combinations cross-reacted with primary antibodies they were not meant to target. First, for our three target antigens (DBH, pERK1/2, ChAT), we used primary antibodies produced in host organisms that were far apart from one another in taxonomic relatedness (mouse, rabbit, and goat; respectively) so cross-reactivity would be minimized. Second, we employed affinity-purified secondary antibodies that were preadsorbed by the supplier against the sera of the other host organisms of secondaries used in our cocktail (*i.e.*, organism cross-adsorbed secondary antibodies).

Footnotes *f–h* of **Table 1** provide details on these cross-adsorption screens. For example, the donkey anti-goat IgG we used was cross-adsorbed with sera from mouse, rat and rabbit; among other taxonomic groups (**Table 1**, footnote *h*). Third, we performed control experiments to test for cross-reactivity. For example, tissue reacted with the rabbit anti-pERK1/2 antibody was incubated with an intentionally-mismatched secondary-fluorophore combination (*e.g.,* biotinylated donkey anti-mouse IgG, followed by streptavidin-Alexa 488; or donkey anti-goat IgG conjugated to Alexa 647). For all of these reactions, cross-reactivity was never observed.

*5.2 Evaluating crosstalk under epifluorescence imaging.* At the level of our epifluorescence microscopy and imaging workflow, we performed single-label immunohistochemical experiments of tissues collected from saline- and 2-DG-treated subjects to ascertain whether an individual signal did not bleed through into the other channels. Some pre-existing single-labeled material (supplied by M. Peveto) was also helpful in this regard. We performed this check by examining each signal within its assigned channel as well as in the other two channels we were capturing. No bleed-through was observed for any of the signals in our epifluorescence system.

*5.3 Evaluating crosstalk under confocal imaging.* We also used our core facility's confocal system to photograph the tissues processed for single-label immunohistochemistry (described above in *Section 2.5.5.2*) to check for cross-talk in that system. We performed additional checks by laser-scanning our triple-labeled material under conditions where the automated sequence of stepwise laser-enabled excitations was reconfigured to allow for individual lasers to be deactivated and the empty channels scrutinized for any tell-tale signals of crosstalk resulting from any fluorophores excited by the remaining active laser [8]. In one instance, we found that the solid-state 550 nm laser produced a red-channel signal (phospho-ERK1/2) that bled through in the far-red channel (ChAT). Additional checks ruled out this as bleed-through from excitation crosstalk of the Cy3 and Alexa-647 fluorophores. Rather, the bleed-through was due to emission crosstalk for the filters configured on the microscope itself. Therefore, to eliminate bleed-through caused by this cross-talk within the configured workflow, the sequence for photographing/scanning triple-labeled material was altered so that the 488-nm and 555-nm lasers performed their sequential scans for the first two channels, and the tissue was photographed in the far-red channel separately.

***6. Anatomical observations for determining regions of interest***

*6.1 Determining atlas level 51.* The *locus ceruleus (Wenzel & Wenzel, 1912)* [6] (LC) is a highly-compact cell group of obliquely arranged neurons that lie adjacent to the floor of fourth ventricle (Reil, 1807–1808) [9] (V4f), medial to the midbrain nucleus of trigeminal nerve (>1840) (MEV), and ventromedial to the *parabrachial nucleus (>1840)* (PB); and is situated in the larger *pontine central gray general (Swanson, 2004)* [10] (PCGg). The adjacent cytoarchitecture of the MEV and the myeloarchitecture of the *superior cerebellar peduncle (Procháska, 1800)* [11] (scp), a fiber tract dividing the PB subnuclei, were key landmarks for determining atlas level and accurate comparison of this region between sections of different brains. In rostral portions of the LC, the cell group was observed to tightly wrap around the lateral border of the *fourth ventricle, principal part (Swanson, 2018)* [12] (V4), then to slightly separate from the margin ventrally in more caudal segments (as depicted in atlas level 51).

The overall shape and height of V4 are also useful indicators of atlas level; however, these are less reliable when dorsal-ventral tilt during sectioning is a known issue, which can elongate or shorten the height of ventricular space in the tissue section. Due to a significant distance between the left and right sides of the region, medial–lateral tilt during sectioning can result in atlas assignment asymmetry between the two sides of the section. In some cases, the plane of section prevented the team member from selecting both left and right-side LC regions, since one of them fell before or after level 51. LC cell groups with a regional boundary that closely matched the LC boundary delineation at atlas level 51 were analyzed. LC ROIs that were determined to be more rostral or caudal than atlas level 51 were not considered here.

The team member marked the brightly-fluorescing perikaryon profiles that had a clearly visible nucleus or closely resembled a segment of a neuron that was consistent in appearance with the cellular morphology of the LC [13]. A perikaryon profile was identified as double-labeled if it appeared yellow in the merged-channels image or, if this was not clear due to one channel obscuring the other, then if it also appeared in both single-channel images. A labeled profile was determined to display only pERK1/2 immunoreactivity or only DBH immunoreactivity by confirming the absence of signal in the other channel for that profile.

*6.2 Determining atlas level 67.* Rostrally, towards what would be *BM4.0* levels 64 and 65, the chemoarchitecture of the *nucleus of solitary tract (>1840)* (NTS) and *dorsal motor nucleus of vagus nerve (>1840)* (DMX) overlap and they appear to share a ventromedial border. Further caudally, however, at atlas level 67, the chemo- and cytoarchitecture of the NTS and DMX are more clearly discernible from one another. The V-shaped *floor of fourth ventricle (Reil, 1807–1808)* [9] (V4f) in this area can be a diagnostic fiducial for comparing these dorsomedially-situated structures between tissue sections. The ventricular opening is much more confined at level 68 as the wider space at level 67 narrows caudally into the *central canal (Bellingeri, 1823)* [14] (C). The *hypoglossal nucleus (>1840)* (XII) neighbors the DMX medially, and its large motor neurons could be identified by ChAT immunolabeling. Moreover, the proximity of XII neurons to those of the DMX were indicative of atlas level, since the borders of the XII and DMX are closer at 67 than at more rostral atlas representations.

The DBH-labeled axonal fibers, in contradistinction, appeared to confine themselves primarily within the borders of the *nucleus of solitary tract, medial part (>1840)* (NTSm) and were used in combination with the descending white matter fibers of the *solitary tract (>1840)* (ts), to localize the NTSm when an adjacent Nissl-stained section was not used. A few stray DBH$^+$ fibers and an occasional DBH$^+$ perikaryon profile were observed lateral to the descending ts fibers; note, however, that we did not include the *nucleus of solitary tract, lateral part (>1840)* (NTSl) in our mapping analysis, but general observations of labeling in this structure were recorded. The reviewer marked each brightly-fluorescing small- to moderately-sized perikaryon profile of the NTS that contained a visible nucleus. Less vibrant or slightly out-of-focus profiles were marked by the reviewer if the perikaryon profile was distinguishable or if the labeled elements displayed features of the other clearly-identifiable DBH-labeled perikaryon profiles, such as a similar perikaryal size or long axonal extensions.

*6.3 Determining atlas level 69.* Atlas level 69 was determined, in the main, by the presence of the *area postrema (>1840)* (AP) dorsal to the DMX and NTS. At level 69, the AP extends laterally and is mostly flat along its dorsal border. At this level, the DMX presents as a large, blimp-shaped cell group that decreases in cellular area and vertically flattens in more caudal representations. Proceeding caudally, the AP takes on a rounder shape as it reduces in size and this shape is a reliable visual cue for the determination of a rostral or caudal AP-containing section. Perikaryon profiles were carefully marked along the border of the NTS and AP so as to not include those found in the AP in our analysis of the NTS.

### 7. Counting procedures

*7.1 Rationale for counting methods.* A few points regarding our counting procedures are noted here. First, in keeping with recommendations furnished by Saper [15], we note here our reasons for choosing the particular counting procedures we employed in this study. The goal of our counting efforts was not to provide an unbiased, global estimate of cell population size within any particular brain structure. Nor was it to provide a formal stereological assessment of absolute cell counts or densities. Rather, the goal was to compare the presence of the cellular activation marker, phospho-ERK1/2, in portions of brain regions with a defined rostrocaudal location and a fairly uniformly-comparable set of cytoarchitectonic boundaries between two experimental groups. Thus, we placed deliberate constraints upon our analysis by selecting regions of interest (ROIs) based on cytoarchitectonic criteria. By doing so, we could perform counts on comparable rostrocaudal levels between groups. Thus, the cytoarchitecture – constrained by plane-of-section analysis and Nissl-/chemoarchitecture-based criteria – was the primary informer of the ROI boundaries we used between groups to perform counts.

Second, the counts in our study are, strictly speaking, those of cell profiles, rather than cells themselves, since approaches used here cannot determine whether the total numbers of profiles sampled correspond directly to total cell counts; indeed, the two metrics are rarely equivalent [16]. Nevertheless, given that the ROIs we compared between groups were constrained to similar locations along the anteroposterior axis and, therefore, also had similar cytoarchitectural boundaries, we reasoned that the relative comparison between median profile counts (and not median cell counts *per se*) would still accurately reveal differences in cellular activation between treatments. To this end, we also corrected for oversampling errors in our profile counts, as described in *Section*

*2.7.4.2.*

*7.2 Abercrombie correction for oversampling.* To perform this correction, we made two measurements: the cell diameter and section thickness. To measure cell diameter, two tissue sections representing two subjects from each treatment group were randomly selected for cell diameter estimation. As described in *Section 2.6.2*, images from these sections were captured using a ×20-magnification objective lens (Plan-Apochromat; N.A., 0.8) on our Zeiss AxioImager M.2 microscope and were examined within Volocity software. The *Measurements* tool within Volocity was used to measure cell type-specific profiles. Two perpendicular lines were drawn across the long and short diameters of each sampled perikaryon profile and the total line lengths (in µm) were recorded in a spreadsheet. Ten randomly-selected cell profiles were measured on each section and the average of the longer diameter of these cell profiles was used as an estimation of cell diameter. The mean perikaryal lengths were calculated for the $DBH^+$ profiles of the LC and NTS, the non-DBH-expressing phospho-ERK1/$2^+$ profiles in the NTS, and the $ChAT^+$ profiles in the DMX.

Second, we used the ×100-magnification objective lens (U Plan S-Apo, oil; N.A., 1.40; working distance, 0.13 mm) on our Olympus BX63 system (described in *Section 2.6.1*) to determine the average thickness of our tissue sections after immunohistochemical processing was completed. To perform this measurement, we opened a live window in cellSense™ Dimension software and, under epifluorescence illumination, opened the fluorescence channel coding for DBH-immunoreactivity (either *red* or *green*, depending on the reaction sets summarized in **Table 1**). A group of $DBH^+$ cells was brought into focus and the *z*-axis positions of the stage (in µm) provided by the software for the top- and bottom-most focal planes containing in-focus neurons were recorded and averaged between groups. The optical thickness of six tissue sections for each subject (n = 11) were recorded in this manner. The average section thickness measures for each subject were pooled and averaged by treatment group (saline: n=5 subjects; 2-DG: n=6 subjects).

With the above two measures now in hand, a cell-type correction factor was calculated for each ROI based on Abercrombie's formula [17]:

$$P= A(M/(L+M))$$

where $P$ = the corrected profile counts, $A$ = the raw profile counts, $M$ = section thickness (in µm), and $L$ = the average length of the perikaryon profile (in µm). The correction factors for each ROI were calculated for each subject and averaged for each treatment group. Perikaryon profile counts in the LC were corrected by multiplying total perikaryon profile counts with the $DBH^+$ cell-type correction factor. Similarly, perikaryon profile counts of all the cell types in the DMX were corrected by multiplying total perikaryon profile counts with the $ChAT^+$ cell-type correction factor. For the NTS, however, we observed a notable size difference between cell types. For example, at *BM4.0* level 69, the average NTS area estimations for $DBH^+$ and pERK1/$2^+$ single-labeled perikaryon profiles are 146 $mm^2$ and 96 $mm^2$, respectively. Thus, for the Abercrombie correction to be performed accurately, we estimated a weighted correction factor for each cell type based on the relative abundance of each cell type in the NTS. Once all data were corrected for across ROIs and subjects, the corrected perikaryon profile counts were the measures used for statistical analysis and graphical presentation.

### Supplemental Tables and Figures (presented in the order in which they were cited in the Main Results; all numeric citations in Figure captions refer to numbered References in main article)

**Table S1**. Summary of descriptive statistics for LC perikaryon profiles at Atlas Level 51

| Region | Cell phenotype | Saline | | | | 2-DG | | | |
|---|---|---|---|---|---|---|---|---|---|
| | | n | Median | Q1 | Q3 | n | Median | Q1 | Q3 |
| LC (~51) | pERK1/2$^+$ | 5 | 14.5 | 11.5 | 17 | 5 | 33 | 31.5 | 37.5 |
| | DBH$^+$ | 5 | 73 | 73 | 86 | 5 | 97.5 | 92.5 | 97.5 |
| | pERK1/2$^+$ & DBH$^-$ | 5 | 1 | 0.5 | 1 | 5 | 0.5 | 0.5 | 1 |
| | pERK$^-$ & DBH$^+$ | 5 | 57 | 53 | 79 | 5 | 63.5 | 61.5 | 65 |
| | pERK$^+$ & DBH$^+$ | 5 | 13 | 11 | 16 | 5 | 32.5 | 30.5 | 37.5 |

**Table S2**. Summary of non-parametric test statistics and effect sizes for LC perikaryon profiles at Atlas Level 51

| Region | Cell phenotype | Sample size | | Wilcoxon Rank-Sum test | | | Wilcoxon effect size | |
|---|---|---|---|---|---|---|---|---|
| | | Saline | 2-DG | Test statistic | *p*-value | FDR-adjusted *p*-value | Effect size | Magnitude* |
| LC (~51) | pERK1/2$^+$ | 5 | 5 | 25 | **0.01** | **0.03** | **0.83** | large |
| | DBH$^+$ | 5 | 5 | 21 | 0.09 | 0.22 | 0.56 | large |
| | pERK1/2$^+$ & DBH$^-$ | 5 | 5 | 6.5 | 0.24 | 0.45 | 0.40 | moderate |
| | pERK1/2$^-$ & DBH$^+$ | 5 | 5 | 14 | 0.83 | 0.45 | 0.10 | small |
| | pERK1/2$^+$ & DBH$^+$ | 5 | 5 | 25 | **0.01** | **0.03** | **0.83** | large |

* 0.1 to < 0.3 – small, 0.3 to < 0.5 – moderate, and > or = 0.5 – large

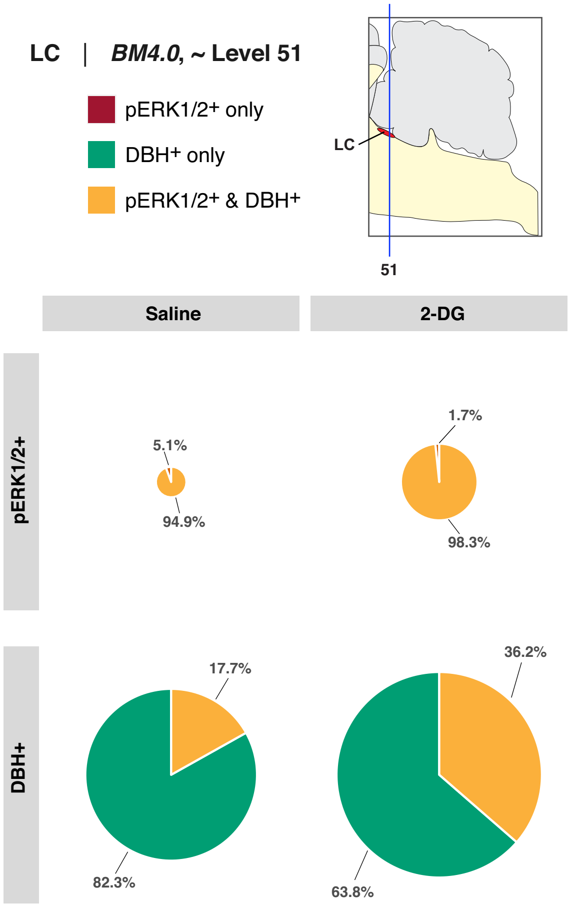


**Figure S1 (*left*).** Relative percent differences of perikaryon phenotypes in *locus ceruleus (Wenzel & Wenzel, 1912)* [83] (LC) region in saline- and 2-DG-treated subjects. Relative percentages of single- and double-labeled perikaryon profiles were calculated for pERK1/2+ (*top row*) and DBH+ (*bottom row*) cell types in each treatment group and displayed as pie charts. Charts are paneled with treatment group as column and perikarya type as row. The size of each pie chart was scaled according to its corresponding total Abercrombie-corrected [100] perikaryon profile number. For example, the 2-DG group pERK1/2+ pie chart (*top right panel*) has a larger diameter than the saline group pERK1/2+ pie chart (*top left panel*), corresponding to the Abercrombie-corrected perikaryon profile numbers presented in **Figure 4**. Note the similarity in the yellow-colored portion of the pie, in terms of relative proportions to the whole, in the top row between treatment groups.

**Table S3**. Summary of descriptive statistics for NTS and DMX perikaryon profiles at Atlas Level 67

| Region | Perikarya phenotype | Saline | | | | 2-DG | | | |
|---|---|---|---|---|---|---|---|---|---|
| | | n | Median | Q1 | Q3 | n | Median | Q1 | Q3 |
| NTS (~67) | pERK1/2$^+$ | 6 | 1.7 | 1.2 | 2.5 | 5 | 6.1 | 6.1 | 7.2 |
| | DBH$^+$ | 6 | 2.1 | 1.6 | 2.3 | 5 | 2.9 | 2.7 | 3.9 |
| | pERK1/2$^+$ & DBH$^-$ | 6 | 1.6 | 0.9 | 2.0 | 5 | 4.2 | 4.0 | 5.5 |
| | pERK1/2$^-$ & DBH$^+$ | 6 | 1.7 | 1.3 | 2.3 | 5 | 1.0 | 0.8 | 2.1 |
| | pERK1/2$^+$ & DBH$^+$ | 6 | 0.2 | 0.2 | 0.3 | 5 | 1.9 | 1.7 | 2.1 |
| DMX (~67) | pERK1/2$^+$ | 6 | 5.4 | 3.3 | 6.1 | 5 | 6.1 | 4.5 | 8.7 |
| | ChAT$^+$ | 6 | 14.9 | 13.9 | 15.6 | 5 | 15.2 | 10.7 | 15.4 |
| | DBH$^+$ | 6 | 0.3 | 0.2 | 0.3 | 5 | 0.9 | 0.7 | 1.0 |
| | pERK1/2$^+$, ChAT$^-$, & DBH$^-$ | 6 | 0 | 0 | 0 | 5 | 0 | 0 | 0 |
| | pERK1/2$^-$, ChAT$^+$, & DBH$^-$ | 6 | 9.4 | 8.1 | 10.7 | 5 | 6.5 | 5.2 | 8.5 |
| | pERK1/2$^-$, ChAT$^-$, & DBH$^+$ | 6 | 0 | 0 | 0 | 5 | 0 | 0 | 0 |
| | pERK1/2$^+$, ChAT$^+$ | 6 | 5.4 | 3.3 | 6.1 | 5 | 6.1 | 4.5 | 8.7 |
| | pERK1/2$^+$, DBH$^+$ | 6 | 0 | 0 | 0 | 5 | 0 | 0 | 0 |
| | ChAT$^+$, DBH$^+$ | 6 | 0.3 | 0.2 | 0.3 | 5 | 0.9 | 0.7 | 1.0 |
| | pERK1/2$^+$, ChAT$^+$, & DBH$^-$ | 6 | 5.2 | 3.3 | 6.0 | 5 | 6.1 | 4.4 | 8.6 |
| | pERK1/2$^-$, ChAT$^+$, & DBH$^+$ | 6 | 0.2 | 0.04 | 0.2 | 5 | 0.9 | 0.7 | 0.9 |
| | pERK1/2$^+$, ChAT$^-$, & DBH$^+$ | 6 | 0 | 0 | 0 | 5 | 0 | 0 | 0 |
| | pERK1/2$^+$, ChAT$^+$, & DBH$^+$ | 6 | 0.2 | 0.03 | 0.2 | 5 | 0.2 | 0 | 0.2 |

**Table S4**. Summary of non-parametric test statistics and effect sizes for NTS and DMX perikaryon profiles at Atlas Level 67

| Region | Perikarya phenotype | Sample size | | Wilcoxon Rank-Sum test | | | Wilcoxon effect size | |
|---|---|---|---|---|---|---|---|---|
| | | Saline | 2-DG | Test statistic | *p*-value | FDR-adjusted *p*-value | Effect size | Magnitude* |
| NTS (~ 67) | pERK1/2$^+$ | 6 | 5 | 30 | **<0.01** | **0.02** | 0.83 | large |
| | DBH$^+$ | 6 | 5 | 23 | 0.17 | 0.38 | 0.44 | moderate |
| | pERK1/2$^+$ & DBH$^-$ | 6 | 5 | 30 | **<0.01** | **0.02** | 0.83 | large |
| | pERK1/2$^-$ & DBH$^+$ | 6 | 5 | 10 | 0.41 | 0.67 | 0.28 | small |
| | pERK1/2$^+$ & DBH$^+$ | 6 | 5 | 29 | **0.01** | **0.02** | 0.78 | large |
| DMX (~67) | pERK1/2$^+$ | 6 | 5 | 18.5 | 0.58 | 0.71 | 0.19 | small |
| | ChAT$^+$ | 6 | 5 | 14 | 0.93 | 1.00 | 0.05 | small |
| | DBH$^+$ | 6 | 5 | 25.5 | 0.06 | 0.19 | 0.59 | large |
| | pERK1/2$^+$, ChAT$^-$, & DBH$^-$ | 6 | 5 | NA | NA | NA | NA | NA |
| | pERK1/2$^-$, ChAT$^+$, & DBH$^-$ | 6 | 5 | 4.5 | 0.07 | 0.19 | 0.58 | large |
| | pERK1/2$^-$, ChAT$^-$, & DBH$^+$ | 6 | 5 | NA | NA | NA | NA | NA |
| | pERK1/2$^+$, ChAT$^+$ | 6 | 5 | 18.5 | 0.58 | 0.71 | 0.19 | small |
| | pERK1/2$^+$, DBH$^+$ | 6 | 5 | NA | NA | NA | NA | NA |
| | ChAT$^+$, DBH$^+$ | 6 | 5 | 25.5 | 0.06 | 0.19 | 0.59 | large |
| | pERK1/2$^+$, ChAT$^+$, & DBH$^-$ | 6 | 5 | 18.5 | 0.58 | 0.71 | 0.19 | small |
| | pERK1/2$^-$, ChAT$^+$, & DBH$^+$ | 6 | 5 | 25 | 0.08 | 0.19 | 0.56 | large |
| | pERK1/2$^+$, ChAT$^-$, & DBH$^+$ | 6 | 5 | NA | NA | NA | NA | NA |
| | pERK1/2$^+$, ChAT$^+$, & DBH$^+$ | 6 | 5 | 16 | 0.92 | 1.00 | 0.06 | small |

* 0.1 to < 0.3 – small, 0.3 to < 0.5 – moderate, and > or = 0.5 – large

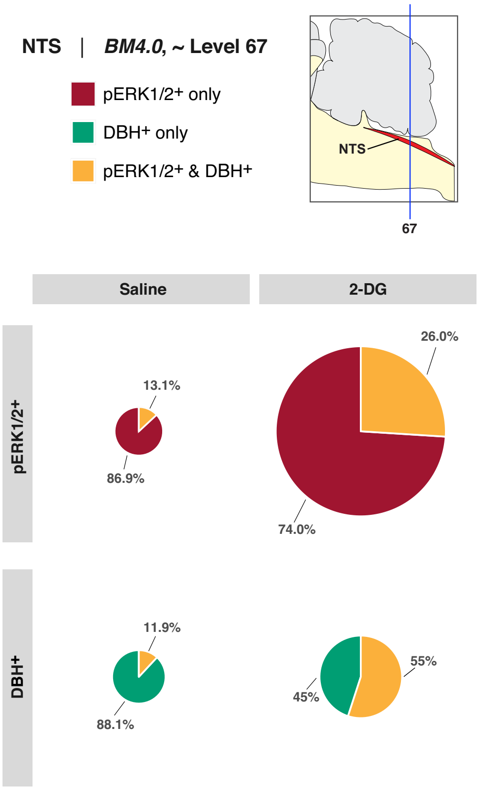


**Figure S2 (*left*).** Relative percent differences of perikaryon phenotypes in *BM4.0* atlas level 67-registered *nucleus of solitary tract (>1840)* (NTS) in saline- and 2-DG-treated subjects. Relative percentages of single- and double-labeled perikaryon profiles were calculated for pERK1/2+ (*top row*) and DBH+ (*bottom row*) cell types in each treatment group and displayed as pie charts. Charts are paneled with treatment group as column and cell type as row. The size of each pie chart was scaled according to its corresponding total Abercrombie-corrected [100] perikaryon profile number. For example, the 2-DG group pERK1/2+ pie chart (*top right panel*) has a larger diameter than the saline group pERK1/2+ pie chart (*top left panel*), corresponding to the Abercrombie-corrected perikaryon profile number presented in **Figure 7**. Note the high percentage of pERK1/2+-only perikaryon profiles in the top row in both saline- and 2-DG-treated rats.

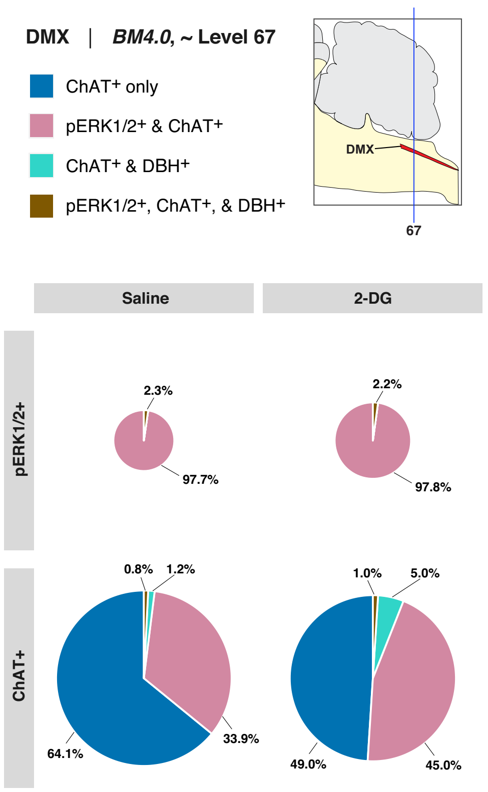


**Figure S3**. Relative percent differences of perikaryon phenotypes in *BM4.0* atlas level 67-registered *dorsal motor nucleus of vagus (>1840)* (DMX) in saline- and 2-DG-treated subjects. Relative percentages of single-, double-, and triple-labeled perikaryon profiles were calculated for pERK1/2+ (*top row*) and ChAT+ (*bottom row*) cell types in each treatment group and displayed as pie charts. Charts are paneled with treatment group as column and cell type as row. The size of each pie chart was scaled according to its corresponding total Abercrombie-corrected [100] perikaryon profile number.

**Table S5**. Summary of descriptive statistics for NTS and DMX perikarya profiles at atlas level 69

| Region | Perikarya phenotype | Saline | | | | 2-DG | | | |
|---|---|---|---|---|---|---|---|---|---|
| | | n | Median | Q1 | Q3 | n | Median | Q1 | Q3 |
| NTS (~69) | pERK1/2$^+$ | 6 | 3.6 | 3.5 | 4.0 | 4 | 8.3 | 8.3 | 9.3 |
| | DBH$^+$ | 6 | 9.7 | 9.3 | 10.4 | 4 | 12.6 | 11.3 | 13.7 |
| | pERK1/2$^+$ & DBH$^-$ | 6 | 2.9 | 2.2 | 3.0 | 4 | 5.7 | 5.3 | 6.1 |
| | pERK1/2$^-$ & DBH$^+$ | 6 | 8.9 | 8.4 | 9.5 | 4 | 8.8 | 8.3 | 9.5 |
| | pERK1/2$^+$ & DBH$^+$ | 6 | 0.8 | 0.7 | 1.1 | 4 | 2.9 | 2.7 | 3.6 |
| DMX (~69) | pERK1/2$^+$ | 6 | 8.8 | 8.0 | 9.6 | 4 | 8.5 | 6.9 | 11.2 |
| | ChAT$^+$ | 6 | 20.9 | 20.3 | 21.4 | 4 | 17.0 | 16.2 | 18.8 |
| | DBH$^+$ | 6 | 0.1 | 0 | 0.1 | 4 | 0 | 0 | 0.04 |
| | pERK1/2$^+$, ChAT$^-$, & DBH$^-$ | 6 | 0 | 0 | 0 | 4 | 0 | 0 | 0 |
| | pERK1/2$^-$, ChAT$^+$, & DBH$^-$ | 6 | 11.9 | 11.5 | 12 | 4 | 8.0 | 7.5 | 8.8 |
| | pERK1/2$^-$, ChAT$^-$, & DBH$^+$ | 6 | 0 | 0 | 0 | 4 | 0 | 0 | 0 |
| | pERK1/2$^+$, ChAT$^+$ | 6 | 8.8 | 8.0 | 9.6 | 4 | 8.5 | 6.9 | 11.2 |
| | pERK1/2$^+$, DBH$^+$ | 6 | 0 | 0 | 0 | 4 | 0 | 0 | 0 |
| | ChAT$^+$, DBH$^+$ | 6 | 0.8 | 0 | 0.1 | 4 | 0 | 0 | 0.04 |
| | pERK1/2$^+$, ChAT$^+$, & DBH$^-$ | 6 | 8.0 | 9.6 | 1.6 | 4 | 6.9 | 11.2 | 4.4 |
| | pERK1/2$^-$, ChAT$^+$, & DBH$^+$ | 6 | 0.8 | 0 | 0.1 | 4 | 0 | 0 | 0.04 |
| | pERK1/2$^+$, ChAT$^-$, & DBH$^+$ | 6 | 0 | 0 | 0 | 4 | 0 | 0 | 0 |
| | pERK1/2$^+$, ChAT$^+$, & DBH$^+$ | 6 | 0 | 0 | 0 | 4 | 0 | 0 | 0 |

**Table S6**. Summary of non-parametric test statistics and effect sizes for NTSm/DMX perikaryon profiles at Atlas Level 69

| Region | Cell phenotype | Sample size | | Wilcoxon Rank-Sum test | | Wilcoxon effect size | |
|---|---|---|---|---|---|---|---|
| | | Saline | 2-DG | Test statistic | *p*-value | Effect size | Magnitude* |
| NTSm (~ 69) | pERK1/2$^{+}$ | 6 | 4 | 24 | **0.01** | 0.82 | large |
| | DBH$^{+}$ | 6 | 4 | 18 | 0.24 | 0.40 | moderate |
| | pERK1/2$^{+}$, DBH$^{-}$ | 6 | 4 | 22.5 | **0.03** | 0.71 | large |
| | pERK1/2$^{-}$, DBH+ | 6 | 4 | 11 | 0.91 | 0.07 | small |
| | pERK1/2$^{+}$, DBH$^{+}$ | 6 | 4 | 24 | **0.01** | 0.82 | large |
| DMX (~69) | pERK1/2$^{+}$ | 6 | 4 | 11.5 | 1.00 | 0.03 | small |
| | ChAT$^{+}$ | 6 | 4 | 8 | 0.45 | 0.27 | small |
| | DBH$^{+}$ | 6 | 4 | 8.5 | 0.46 | 0.27 | small |
| | pERK1/2$^{+}$, ChAT$^{-}$, DBH$^{-}$ | 6 | 4 | NA | NA | NA | NA |
| | pERK1/2$^{-}$, ChAT$^{+}$, DBH$^{-}$ | 6 | 4 | 3 | 0.07 | 0.60 | large |
| | pERK1/2$^{-}$, ChAT$^{-}$, DBH$^{+}$ | 6 | 4 | NA | NA | NA | NA |
| | pERK1/2$^{+}$, ChAT$^{+}$ | 6 | 4 | 11.5 | 1.00 | 0.03 | small |
| | pERK1/2$^{+}$, DBH$^{+}$ | 6 | 4 | NA | NA | NA | NA |
| | ChAT$^{+}$, DBH$^{+}$ | 6 | 4 | 8.5 | 0.46 | 0.27 | small |
| | pERK1/2$^{+}$, ChAT$^{+}$, DBH$^{-}$ | 6 | 4 | 11.5 | 1.00 | 0.03 | small |
| | pERK1/2$^{-}$, ChAT$^{+}$, DBH$^{+}$ | 6 | 4 | 8.5 | 0.46 | 0.27 | small |
| | pERK1/2$^{+}$, ChAT$^{-}$, DBH$^{+}$ | 6 | 4 | NA | NA | NA | NA |
| | pERK1/2$^{+}$, ChAT$^{+}$, DBH$^{+}$ | 6 | 4 | 10 | 0.54 | 0.25 | small |

* 0.1 to < 0.3 – small, 0.3 to < 0.5 – moderate, and > or = 0.5 – large

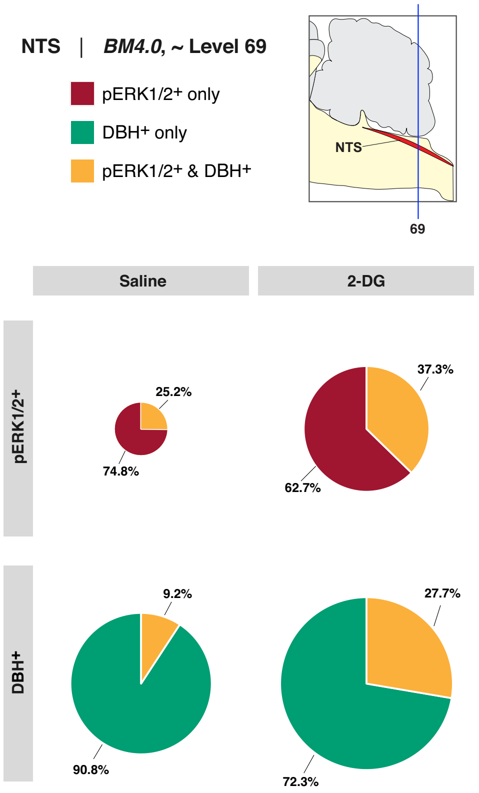


**Figure S4**. Relative percent changes of perikaryon profile phenotypes in *BM4.0* atlas level 69- registered *nucleus of solitary tract (>1840)* (NTS) 15 min after 2-DG administration. Relative percentages of single and double-labeled perikarya profiles were calculated for pERK1/2+ (*top row*) and DBH+ (*bottom row*) perikaryon types in each treatment group and displayed as pie charts. Charts are paneled with treatment group as column and perikarya type as row. The size of each pie chart was scaled according to its corresponding total Abercrombie-corrected [100] perikaryon profile number. For example, the 2-DG group's pERK1/2+ pie chart (*top right panel*) has a larger diameter than the saline group pERK1/2+ pie chart (*top left panel*), corresponding to the Abercrombie-corrected perikaryon profile number presented in **Figure 13**. Notice the high percentage of pERK1/2+ only perikarya in the top row in both saline- and 2-DG-treated rats.

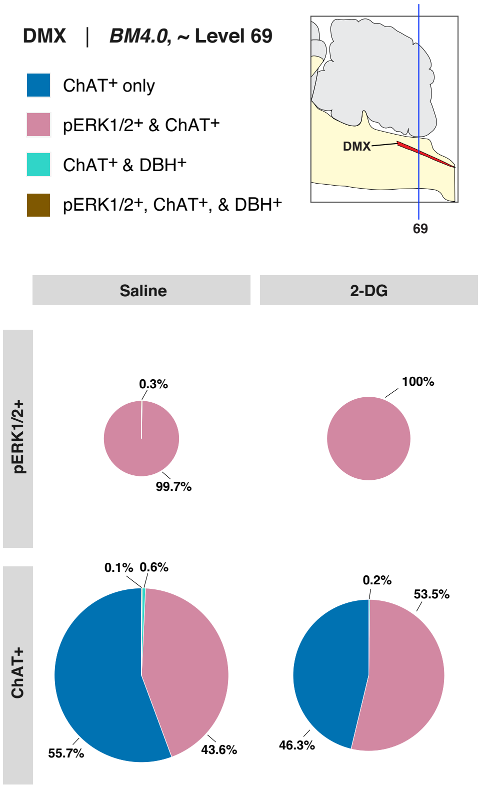


**Figure S5**. Relative percent changes of perikaryon phenotypes in the *BM4.0* atlas level 69-registered *dorsal motor nucleus of vagus nerve (>1840)* (DMX) 15 min after 2-DG administration. Relative percentages of single-, double-, and triple labeled perikarya profiles were calculated for pERK1/2+ (*top row*) and ChAT+ (*bottom row*) perikaryon types in each treatment group and displayed as pie charts. Charts are paneled with treatment group as column and perikaryon type as row. The size of each pie chart was scaled according to its corresponding total Abercrombie-corrected perikaryon profile numbers.
